# Intragenic epistasis in couch potato shapes life history traits and climatic adaptation in *Drosophila melanogaster*

**DOI:** 10.64898/2026.10.09.758124

**Authors:** Emily L. Behrman, Alan O. Bergland, Dmitri A. Petrov, Paul Schmidt

## Abstract

Populations adapting to heterogeneous environments often harbor many alleles that covary with environmental gradients, yet how so many adaptive variants can be maintained and show signatures of local adaptation remains unclear. Epistatic non-additive interactions among alleles may concentrate fitness effects at specific allelic combinations to shape complex traits and facilitate adaptation. We evaluate compositional epistasis among three intronic SNPs in the *couch potato* (*cpo*) gene, each independently clinal along the North American latitudinal gradient, using recombinant outbred populations fixed for distinct three-locus combinations. Flies carrying all three temperate alleles (*cpo*^TTA^) or all three tropical alleles (*cpo*^CGT^) show non-additive increases in traits associated with fitness in their respective environments: dormancy, starvation resistance, and lipid storage in temperate flies and fecundity in tropical flies. Intermediate genotype combinations generally show weaker and negative deviations from additive predictions. Temperate genotypes show reduced *cpo* expression and independently create predicted transcription factor binding sites, suggesting a regulatory basis for the observed phenotypic effects. Consistent with selection shaping haplotype frequencies against a backdrop of ongoing recombination, the temperate and tropical genotypes are enriched above random expectation across the latitudinal cline, with directionally consistent but non-significant seasonal patterns. Our findings demonstrate that non-additive interactions among non-coding variants within a single gene can produce large, pleiotropic fitness effects and shape genotype distributions in natural populations, providing a mechanism by which a small number of regulatory variants can underlie complex climatic adaptation.

## Introduction

A longstanding principle in evolutionary biology is that traits or genotypes that covary along spatial environmental gradients (e.g. latitude, longitude and altitude) are inferred to be involved in climatic adaptation (Endler 1977). Evidence of adaptation is even stronger when parallel patterns are replicated across different gradients. The application of genomic approaches to study natural populations reveals that many species contain a vast number of polymorphisms that correlate with environmental clines (Turner et al. 2008; Yi et al. 2010; Ellison et al. 2011; Hancock et al. 2011; Kolaczkowski et al. 2011; Jeffery et al. 2017; Stankowski et al. 2017; Phifer-Rixey et al. 2018; Westram et al. 2018). The large volume of clinal single nucleotide polymorphisms (SNPs) raises the question of whether individual variants act independently and whether they reflect spatially varying selection or demographic history. Disentangling which clinal SNPs are causal in local adaptation from those whose patterns result from genetic drift, genetic draft, or demographic structure remains a challenge.

One approach to studying which clinal SNPs are truly causal is to examine variation within genes rather than treating each SNP as an independent unit (Durmaz et al. 2019; Betancourt et al. 2021). SNPs within a gene may interact epistatically despite low linkage disequilibrium such that each variant appears independently adaptive in isolation while the true fitness effect is a property of the combination, thus potentially inflating the apparent number of adaptive loci. Additive interactions among loci produce phenotypes equivalent to the sum of each allele’s effect, while epistatic interactions result in phenotypes that deviate from this additive expectation by either exceeding (positive epistasis) or falling below (negative epistasis) the additive prediction (Bateson 1906). Epistasis is increasingly recognized as a major force shaping the course and outcome of adaptive evolution, with consequences for fitness landscapes, evolutionary trajectories, and the maintenance of polymorphism (Bank 2022). Epistasis among existing genetic variants may partially explain the prevalence of SNPs inferred to be adaptive. However, the effect size of individual SNPs and the architecture of local adaptation remain poorly resolved, due to a paucity of comprehensive investigations into the functional significance of segregating polymorphisms in natural populations.

Complex traits and genetic polymorphisms distributed over spatial and temporal gradients that are inferred to be products of natural selection have been studied extensively in *Drosophila melanogaster* (Berry and Kreitman 1993; James and Partridge 1995; Verrelli and Eanes 2000; Mitrovski and Hoffmann 2001; Sezgin et al. 2004; Sgrò et al. 2010). A suite of traits covary predictably with climatic gradients across latitude; flies collected from temperate climates are characterized by higher stress resistance, including reproductive dormancy, cold tolerance, starvation resistance, and sometimes longevity, while flies from warmer climates tend to have higher reproductive output (Capy et al. 1993; De Jong and Bochdanovits 2003; Schmidt et al. 2005; Schmidt and Paaby 2008; Betancourt et al. 2021). Seasonal climatic shifts within temperate *D. melanogaster* populations produce phenotypic patterns paralleling the latitudinal cline with higher stress resistance in the spring after winter selection and higher reproductive output in the autumn after summer population growth (Schmidt and Conde 2006; Behrman et al. 2015; Behrman et al. 2018; Rajpurohit et al. 2018; Behrman et al. 2020; Erickson et al. 2020; Behrman and Schmidt 2022). Population sequencing of *D. melanogaster* across spatial and temporal gradients reveals hundreds to thousands of SNPs associated with climatic adaptation (Turner et al. 2008; Kolaczkowski et al. 2011; Fabian et al. 2012; Bergland et al. 2014; Reinhardt et al. 2014; Bergland et al. 2016; Kapun et al. 2020; Kapun et al. 2021; Machado et al. 2021; Behrman and Schmidt 2022; Yu and Bergland 2022). The numerous SNPs that covary with latitude and season may interact in non-additive ways to produce the complex traits sculpted by natural selection during climatic adaptation. Dissecting the functional significance of intragenic interactions requires direct, gene-specific experimental investigation.

The putative RNA binding protein *couch potato* (*cpo*) segregates for multiple polymorphisms that are strongly clinal and seasonal in wild *D. melanogaster* (Schmidt et al. 2008; Fabian et al. 2012; Bergland et al. 2014; Cogni et al. 2014; Kapun et al. 2021; Machado et al. 2021; Behrman and Schmidt 2022). *cpo* is a major locus determining the frequency of reproductive dormancy essential for overwintering in temperate climates across multiple dipteran species, including *D. melanogaster* (Schmidt et al. 2008), *Culex pipiens* (Zhang and Denlinger 2011), *D. montana* (Kankare et al. 2010), and *D. virilis* (Salminen et al. 2015). Dormancy is correlated with differential expression of *cpo*, in some cases in a tissue-specific or dormancy stage-specific pattern (Schmidt et al. 2008; Zhang and Denlinger 2011; Salminen et al. 2015; Zhao et al. 2015). The molecular mechanisms by which natural variants in *cpo* alter life history traits remain unclear, but the pleiotropic effects are predicted to reflect the post-transcriptional role of an RNA binding protein in gene expression regulation. A simple two-SNP genetic architecture at *cpo* (Ala/Val347 and 48034(A/T)) correlates with the latitudinal cline in reproductive dormancy propensity in North American *D. melanogaster* (Schmidt et al. 2008); however, only the Ala/Val347 variant is associated with the weaker dormancy cline in Europe (Zonato et al. 2016), and the role of *cpo* polymorphisms in the Australian dormancy cline could not be confirmed (Lee et al. 2011). These geographic inconsistencies suggest that reproductive dormancy may have a more complex genetic architecture than individual SNPs can capture. The role of *cpo* in climatic adaptation may depend on genetic background and intragenic context.

Here, we identify three highly clinal SNPs in *cpo* (dm6 positions 3R:17964408, 3R:17965558, and 3R:17967866) that are in low linkage disequilibrium with one another and thus appear to act as independent variants, yet they interact non-additively to produce phenotypes that deviate substantially from additive predictions. We test whether these three SNPs have independent functional effects and whether they interact non-additively to produce more extreme trait values that could increase fitness in local environments. If these alleles are functional, we predict that flies carrying the three temperate alleles (*cpo^TTA^*) will show higher performance for traits associated with temperate climate adaptation—including dormancy, stress resistance, lipid storage, and longevity—while flies carrying the three tropical alleles (*cpo^CGT^*) will show higher fecundity, consistent with tropical climate adaptation. We further predict that if these combinations are maintained by selection, the intact haplotypes will occur at higher than expected frequencies given the individual allele frequencies in natural populations, with clinal and seasonal patterns mirroring those of the individual SNPs. We find that the intact temperate and tropical three-locus combinations show non-additive fitness advantages in their respective climatic environments consistent with our predictions and these combinations are maintained at elevated frequences in wild populations.

## Results

### Independently clinal and seasonal loci in *cpo*

Loci in *cpo* potentially involved in climatic adaptation were identified by allele frequencies that covaried with latitude along the east coast of North America. Forty-five SNPs in *cpo* had strongly clinal q-values (CQ) less than 10^-4^ (Figure 1A; Table S1; (Bergland et al. 2014). The clinal SNPs were distributed across the gene and located primarily in non-coding regions. Linkage disequilibrium among the clinal SNPs was examined in the *Drosophila* Genetics Reference Panel (DGRP; (Mackay et al. 2012)): 21 SNPs formed a linkage disequilibrium (D’) block between 3R:17940310 and 3R:17963448 (*D. melanogaster* reference genome v.6), but the remaining SNPs had low average pairwise D’ and appeared to assort independently (Figure 1B).

**Figure 1.**
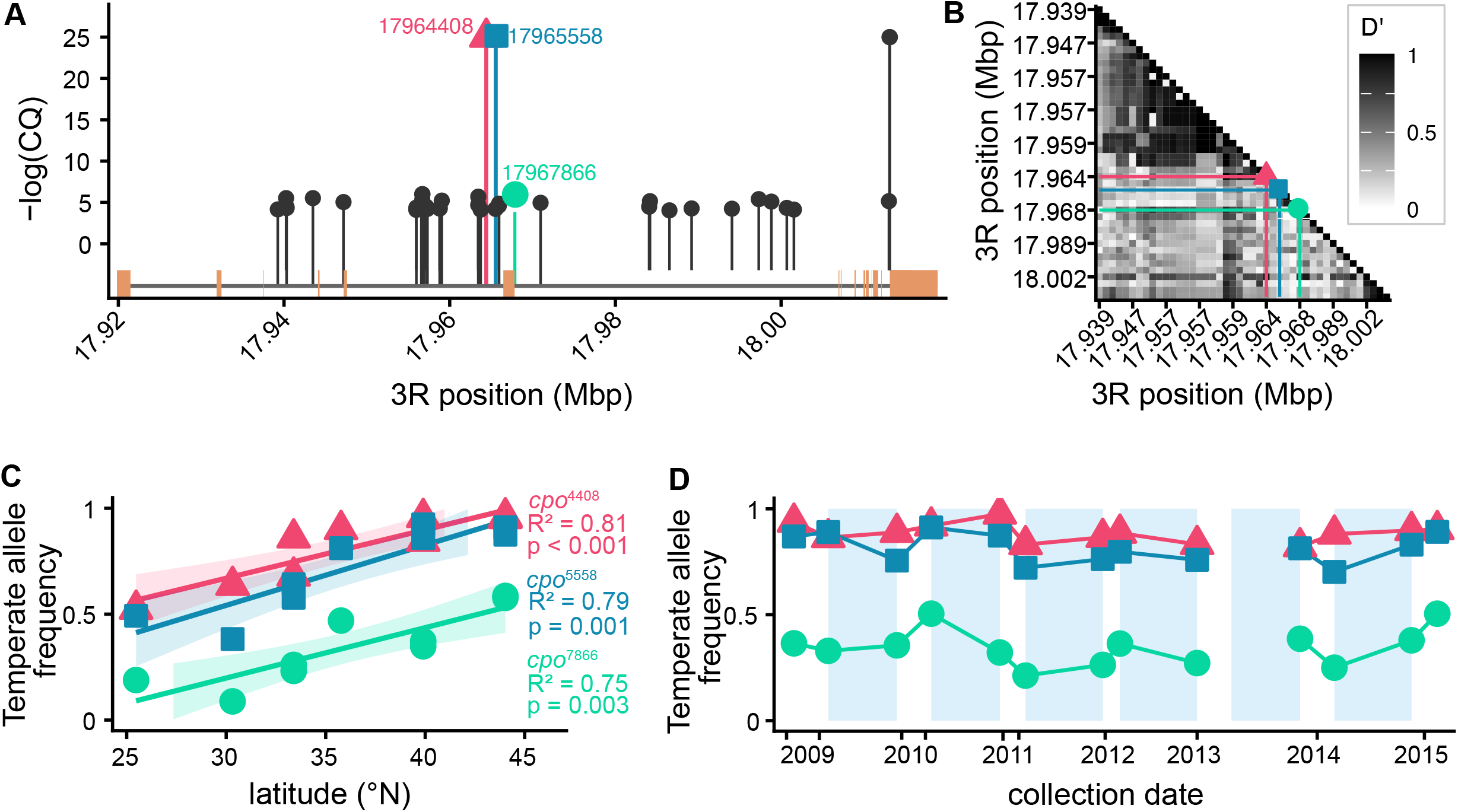
Clinal SNPs in *couch potato* (*cpo*) along the east coast of North America. (A) Clinal Q value (CQ) of SNPs in *cpo* with a CQ <10^-4^ as a function of position in the gene and focal SNPs are highlighted with SNP-specific colored shapes: 3R:17964408 (pink triangle), 3R:17965558 (blue square), 3R:17967866 (green circle). The genic structure of *cpo* is along the X axis. (B) Pairwise linkage disequilibrium (D’) among clinal SNPs with a cutoff of CQ <10^-4^ based on the Drosophila Genetic Reference Panel. (C) Variation in allele frequency of focal *cpo* SNPs as a function of latitude. (D) Variation in allele frequency of focal cpo SNPs as a function of seasonal time in the spring and autumn across six consecutive years at Linvilla Orchards, Media, PA. Winter is highlighted in light blue boxes.

We assessed non-additive interactions among the SNP associated with female reproductive dormancy (Schmidt et al. 2008), 3R:17967866 (*cpo^7866^*, CQ = 2×10^-5^), and two upstream, highly clinal SNPs with CQ < 10^-6^: 3R:17964408 (*cpo^4408^*) and 3R:17965558 (*cpo^5558^*, Figure 1B). These three focal SNPs spanned approximately 3kb and together formed eight genotype combinations that all occurred in natural populations (Mackay et al. 2012). D’ in the DGRP was low between *cpo*^4408^ and *cpo^5558^*(D’=0.186) and between *cpo^5558^* and *cpo^7866^* (D’=0.123), though *cpo^4408^* and *cpo^7866^* showed moderate co-variation (D’=0.514; Figure 1C).

### *cpo* genotypes had pleiotropic and non-additive fitness effects

First, we assessed the effects of individual loci on climatic adaptation using recombinant outbred populations (ROPs) derived from DGRP inbred lines (Mackay et al. 2012), each fixed for one combination of the three focal SNPs in a heterogeneous genomic background (Figure S1; (Paaby et al. 2014)).

We evaluated whether alternative temperate or tropical alleles at each locus independently contributed to trait values by calculating the average trait value among all ROPs containing each alternative allele at a single locus. Some individual loci had significant effects on dormancy, starvation survival, lipid mass, lifespan, or development time (Table S3). However, the directionality of effects did not consistently correspond with allele source, as temperate alleles did not always confer higher performance for traits associated with temperate climate adaptation and vice versa.

Next, we assessed the roles of the temperate (*cpo^TTA^*) and tropical (*cpo^CGT^*) genotypes in environmental adaptation using ROPs fixed for all three alleles from each respective population. The relative performance of the two genotypes was largely consistent with predictions based on local adaptation (Figure 2, Table S4). Flies carrying all three temperate alleles showed higher performance for stress traits associated with temperate survival, including reproductive dormancy (χ^2^(1)=10.01, p=0.0015), starvation resistance (t(473)=-6.97, p=1.1×10^-11^), and lipid content (t(80)=-2.35, p=0.021), compared to flies carrying all three tropical alleles. Conversely, flies carrying all three tropical alleles had higher reproductive output than temperate flies (t(17)=2.54, p=0.021), consistent with adaptation to tropical environments. Counter to climate-based predictions, temperate flies had significantly shorter lifespan than tropical flies (t(1524)=12.7, p=2.2×10^-16^), a pattern that may reflect a life history tradeoff between dormancy investment and longevity under benign laboratory conditions. Thermal tolerance did not differ significantly between genotypes.

**Figure 2.**
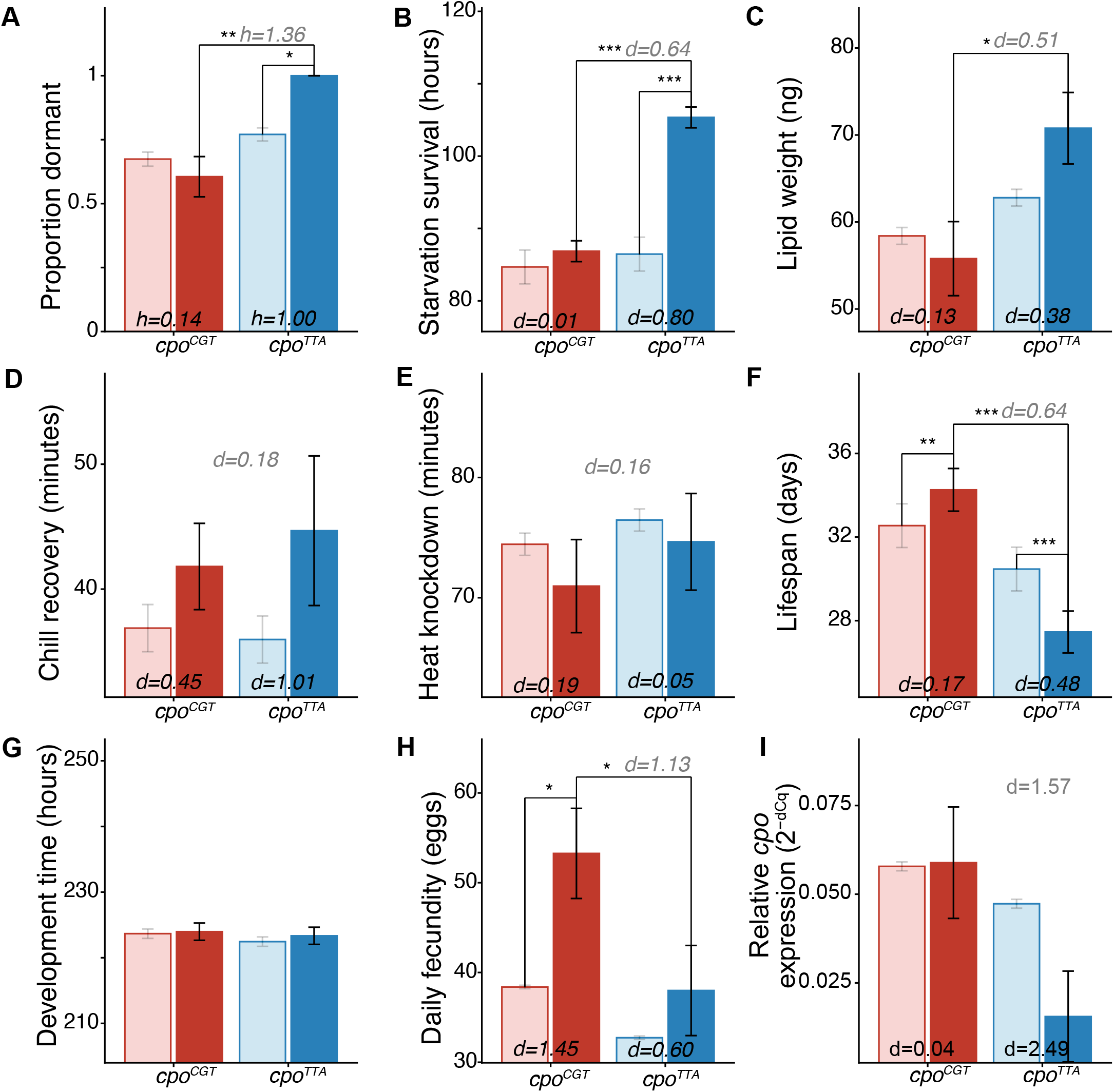
Observed versus predicted trait values in recombinant outbred populations (ROPs) fixed for the tropical (*cpo*^CGT^) and temperate (*cpo*^TTA^) three-locus combinations. For each trait, light bars show the additive null prediction (mean of marginal single-locus effects); dark bars show the observed value for that combination. Significance brackets indicate: within-genotype deviation from the additive prediction (CGT and TTA separately) and the cross-genotype comparison (CGT vs. TTA, observed values), assessed by t-test (*p<0.05, **p<0.01, ***p<0.001). Effect sizes are Cohen’s d for continuous traits and Cohen’s h for the binomial proportions. (A) Proportion dormant. (B) Starvation survival. (C) Lipid weight. (D) Chill coma recovery. (E) Heat knockdown time. (F) Lifespan. (G) Development time. (H) Daily fecundity. (I) Relative *cpo* expression (2-ΔCq). Error bars show ±1 SE. Bottle/vial clustering was accounted for as a random effect where applicable.

We then examined whether temperate and tropical trait values were additive or whether non-additive interactions among loci produced phenotypes deviating from additive expectations. A null predicted value was calculated for each genotype assuming each locus contributed additively, computed as the average trait value across ROPs containing a mixture of temperate and tropical alleles at each of the three loci separately (Figure S1). Epistasis was inferred from deviations between this null prediction and the measured trait value of the corresponding ROP. Both temperate and tropical genotypes showed positive epistasis for traits associated with higher fitness in their respective climatic environments (Figure S1, Table S4). Temperate flies showed higher stress resilience than additive predictions, with higher reproductive dormancy (one-sample proportion test, χ^2^(1)=5.61 p=0.0178) and longer starvation survival (t(9)=-12.07, p=7.33×10^-7^). Tropical flies laid more eggs than predicted (t(9)=3.19, p=0.0102). A complementary likelihood-ratio test comparing the additive model to the full genotype-combination model, which tests for non-additive effects across all seven genotypes rather than at specific vertices, converged with the vertex-specific results for most traits(dormancy: χ^2^(3)=10.12 p=0.018, starvation: χ^2^(3)=94.94 p=1.9×10^-20^, development: χ^2^(3)=24.41 p=2.1×10^-5^, fecundity: F(3,54)=8.03 p=0.0048), with the exception of lifespan (χ^2^(3)=4.79, p=0.188, Table S6). Together, these results show that non-additive fitness effects are broadly concentrated at the temperate and tropical endpoints rather than the intermediate genotypes. Flies carrying intermediate genotypes that contained a mixture of temperate and tropical alleles primarily matched additive predictions or showed negative epistasis, with trait values below additive expectations (Figure 3).

**Figure 3.**
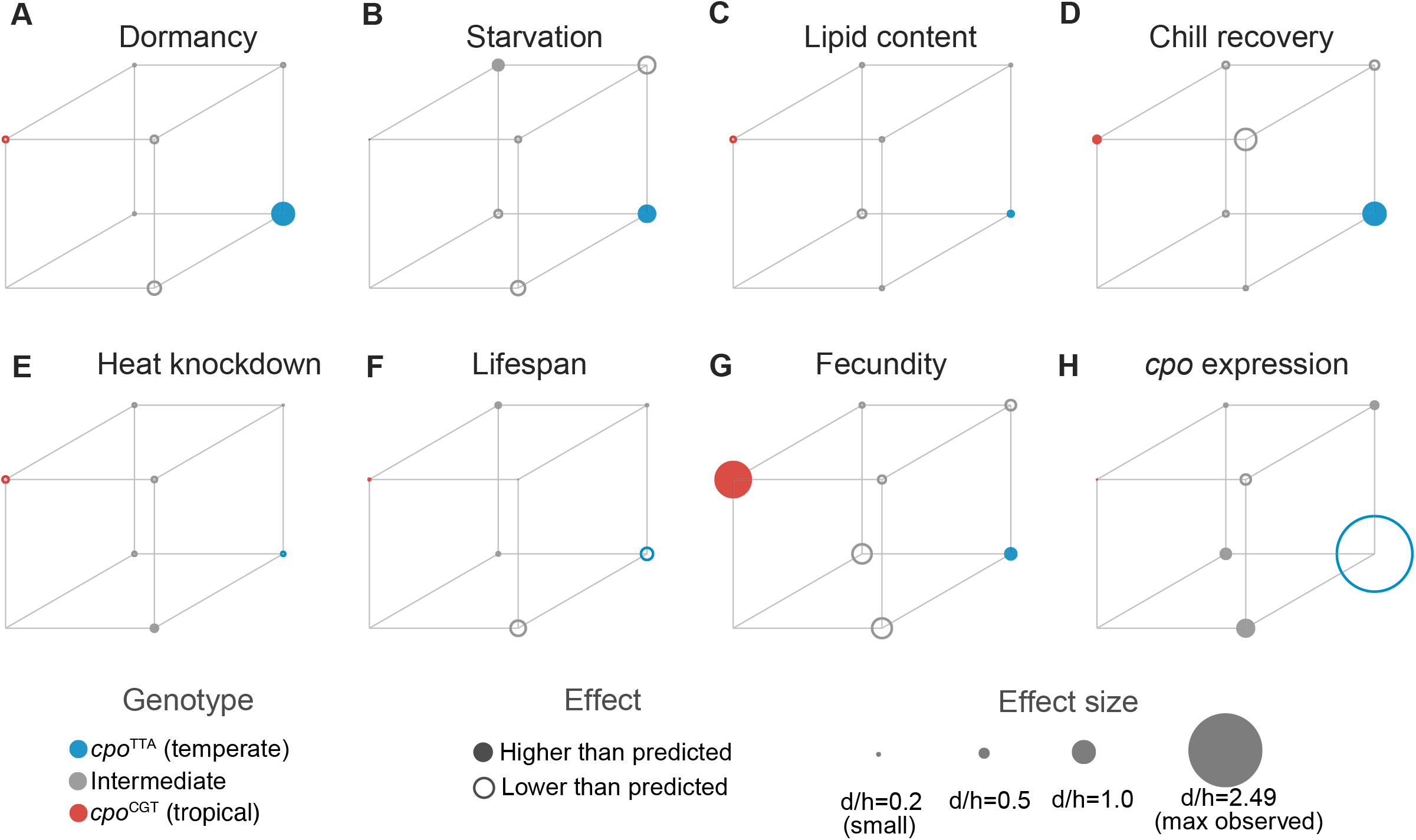
Fitness landscape of compositional epistasis across the seven three-locus *couch potato* (*cpo*) genotype combinations, for eight fitness-related traits. Each cube vertex represents one genotype combination; vertex position follows the three focal SNPs (*cpo*^4408^, *cpo*^5558^, *cpo*^7866^). Marker fill indicates direction of deviation from the additive null prediction (filled = higher than predicted, open = lower than predicted); marker size is proportional to the magnitude of that deviation, scaled consistently across all panels. The *cpo*^CGA^ combination (front, left, bottom corner, blank vertex in all panels) was too rare in the DGRP to establish a recombinant outbred population and is not represented. Colors indicate the *cpo*^TTA^ (temperate, blue), *cpo*^CGT^ (tropical, red), and intermediate (grey) genotypes. (A) Proportion dormant. (B) Starvation survival. (C) Lipid weight. (D) Chill coma recovery. (E) Heat knockdown time. (F) Lifespan. (G) Daily fecundity. (H) Relative *cpo* expression.

We replicated the epistatic effects observed in ROPs across independent genetic backgrounds by assaying chill recovery, starvation resistance, and heat knockdown in Pennsylvania inbred lines fixed for each of the eight *cpo* genotype combinations (Behrman et al. 2018), Figure S2). Consistent with the ROP results, temperate genotype *cpo^TTA^* flies survived starvation significantly longer than predicted by the additive null (p=0.0001) and longer than tropical genotype *cpo^CGT^* flies (p=0.016). Tropical *cpo^CGT^* flies showed a marginally non-significant excess of starvation survival over the additive prediction (p=0.055), and recovered from chill significantly faster than temperate flies (p=0.007). Directional patterns were concordant across both experimental systems, although effect sizes were smaller in the inbred lines, consistent with reduced statistical power in an isogenic background.

### The fitness landscape of *cpo* genotypes is concentrated at temperate and tropical endpoints

Visualizing the deviation from additive predictions across all the *cpo* genotype combinations reveals that non-additive effects are predominantly concentrated at the temperate *cpo*^TTA^ and tropical *cpo*^CGT^ vertices of the genotypic landscape, with intermediate genotypes generally showing weaker deviations (Figure 3, Figure S2). If epistasis arose through a stepwise adaptive walk, we would expect fitness to increase continuously along a path of sequential single-locus substitutions from one endpoint genotype to the other with each intermediate genotype along that path showing a correspondingly intermediate deviation from the additive prediction rather than the sharp concentration of positive deviation at the two endpoints alone that we observe (Weinreich et al. 2005; Weinreich et al. 2006). Instead, the fitness benefits appear to emerge specifically from the intact three-allele combinations. This pattern was clearest for fecundity, where both endpoint genotypes showed positive deviation while all five intermediate genotypes showed negative deviation, and largely held for reproductive dormancy. Starvation resistance and lifespan showed partial exceptions: one intermediate genotype (*cpo*^CTT^) showed a starvation deviation comparable in magnitude to *cpo*^TTA^, and one intermediate genotype (*cpo*^TGA^) showed a lifespan deviation exceeding that of either endpoint. The overall concentration of epistasis at the endpoints rather than uniformly across intermediate genotypes remains inconsistent with a simple stepwise adaptive walk and is more consistent with fitness peaks accessible from standing genetic variation (Barrett and Schluter 2008; Bank 2022; Diaz-Colunga et al. 2023), though the *cpo*^CTT^ and *cpo*^TGA^ exceptions suggest the underlying fitness landscape is not perfectly two-peaked and may include additional local structure not captured by the temperate/tropical endpoint framework. This pattern is also consistent with the low but non-random linkage disequilibrium among the focal SNPs in natural populations, where selection appears to maintain the intact combinations despite ongoing recombination.

### Genotype-specific *cpo* expression

We hypothesized that the intronic focal SNPs may affect expression of *cpo* transcripts, contributing to the widespread phenotypic differences among genotypes. Relative whole-body *cpo* mRNA expression in temperate genotype females was approximately 3.8-fold lower than in tropical genotype females (Figure 2; Table S4). The additive null model predicted lower *cpo* expression in temperate than tropical flies, and while temperate flies showed a large reduction in expression relative to the additive prediction, this deviation did not reach significance (p=0.093, Cohen’s d=2.49), likely reflecting low statistical power rather than a true null effect. Tropical flies had expression levels equivalent to the additive prediction.

### Focal SNPs are predicted to create ecdysone receptor and NRF2 binding sites

Given their intronic location and the observed genotype-specific expression differences, we asked whether the focal SNPs fall within known *Drosophila* transcription factor binding motifs. Across all significant motif disruptions, the temperate allele produced a stronger predicted binding score than the tropical allele in 22 of 28 cases (78.6%; Figure 4A), establishing a general directional bias.

**Figure 4.**
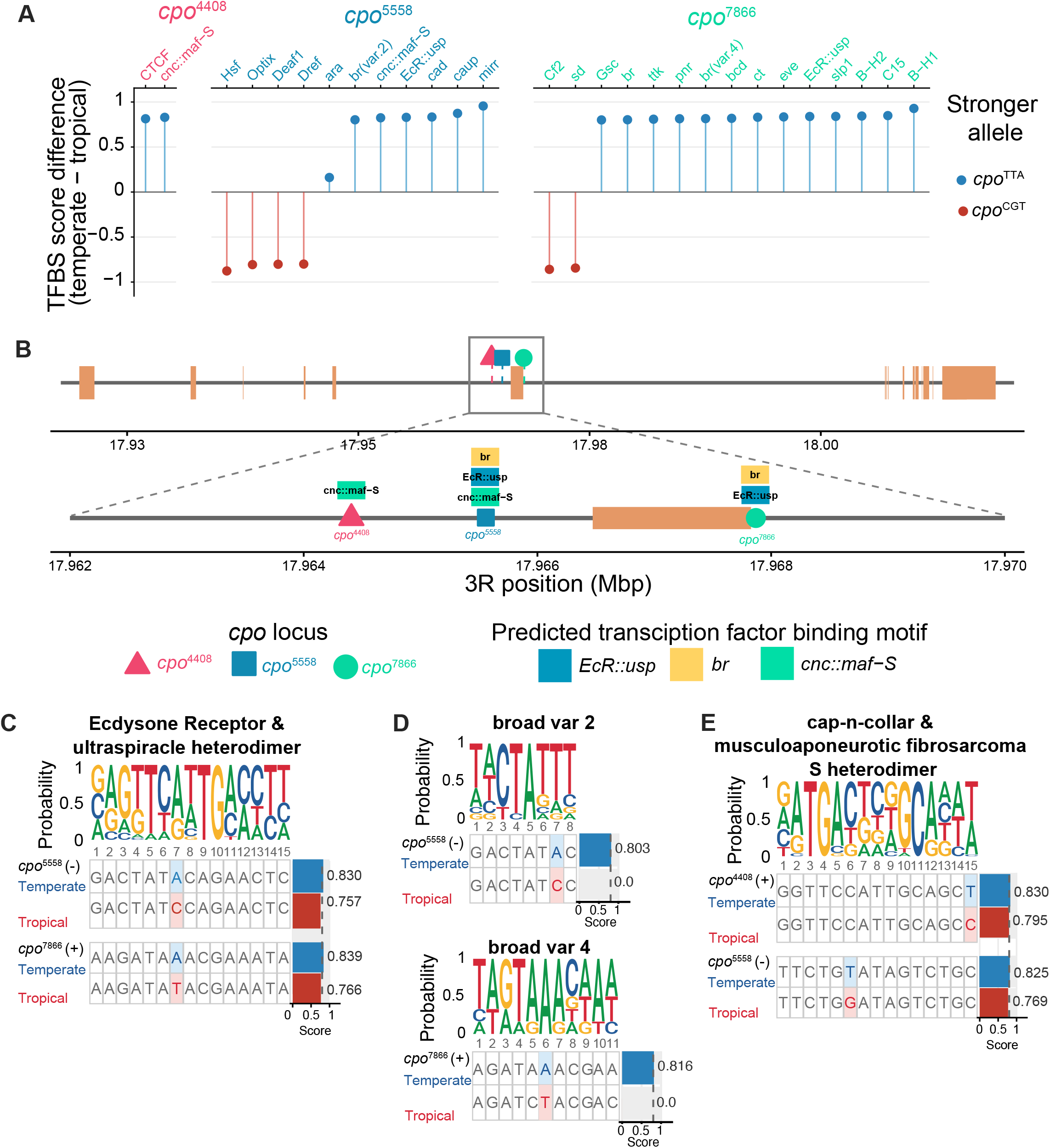
Predicted transcription factor binding site (TFBS) disruption at the three focal cpo SNPs. (A) Score difference (temperate − tropical) for all 28 significant motif disruptions (|Δscore| > 0.1) across the three loci, colored by which allele binds more strongly (blue = temperate stronger; red = tropical stronger). The temperate allele is stronger in 22 of 28 cases (78.6%). (B) *couch potato* (*cpo*) gene structure with the three focal SNPs (triangle/square/circle) and their disrupted motifs, color-coded by transcription factor. (C–E) Sequence logos and allele-specific binding comparisons for the three regulatory path-ways: (C) *Ecdysone receptor/Ultraspiracle* (*EcR::usp*) at *cpo*5558 and *cpo*7866; (D) *Broad* (*br*) isoforms *br*(var.2) and *br*(var.4); (E) *Cap-n-collar/musculoaponeurotic fibrosarcoma* (*cnc::maf-S*) at *cpo*4408 and *cpo*5558. Top row of each pair shows the temperate allele (blue), bottom shows tropical (red); the focal SNP position is highlighted, and strand orientation is noted beside each locus label. Bars show relative binding score (0–1); dashed line marks the 80% threshold for a called site.

The most biologically compelling result was the independent creation of predicted ecdysone receptor (*EcR::usp*) binding sites by the temperate alleles at two of the three focal SNPs (Figure 4B). At *cpo*^5558^, the temperate allele (T) produced an *EcR::usp* motif score of 0.830, exceeding the 80% binding threshold, while the tropical allele (G) scored 0.757, below threshold (site sequence: GACTATACAGAACTC vs. GACTATCCAGAACTC; Figure 4C). At *cpo*^7866^, the temperate allele (A) similarly crossed the threshold (score=0.839) while the tropical allele (T) did not (score=0.766; site sequence: AAGATAAACGAAATA vs. AAGATATACGAAATA; Figure 4C). Strikingly, both SNPs independently altered the same position — position 7 — of the 15-base *EcR::usp* motif, the most information-rich position in the binding site (2.0 bits). The probability of two independent SNPs both targeting position 7 by chance is (1/15)^2^ = 0.0044.

The temperate alleles at *cpo*^4408^ and *cpo*^5558^ each affected predicted cnc::maf-S binding, the *Drosophila* ortholog of the NRF2 antioxidant response transcription factor (Figure 4B). At *cpo*^4408^, the temperate allele (T) retained a *cnc::maf-S* site at motif position 15 (score=0.830), while the tropical allele (C) fell below threshold (score=0.795; Figure 4D). At *cpo*^5558^, the temperate allele created a stronger *cnc::maf-S* site (score=0.825) than the tropical allele (score=0.769; Figure 4D).

Focal SNPs also created predicted binding sites for *Broad* complex (*br*) isoforms (Figure 4B) — ecdysone-responsive zinc finger transcription factors downstream of *EcR::usp* signaling. At *cpo*^5558^, the temperate allele created a predicted *br*(var.2) binding site absent in the tropical allele (score=0.803 vs. 0.000; Figure 4E). At *cpo*^7866^, the temperate allele created a predicted *br*(var.4) binding site absent in the tropical allele (score=0.816 vs. 0.000; Figure 4E). *Broad* complex binding sites (any isoform) were created by the temperate alleles at 2 of 3 focal SNPs, compared to 18 of 71 non-focal SNPs (Fisher’s exact p=0.176). This lack of significance likely reflects the comparatively high background rate of *Broad* motif disruption across the region (25% of non-focal SNPs) relative to EcR::usp (8%) or cnc::maf-S (4%), consistent with *Broad*’s more degenerate binding consensus. Nonetheless, the convergent creation of *Broad* binding alongside *EcR::usp* sites is consistent with the temperate haplotype establishing a coordinated ecdysone-responsive regulatory module.

To assess whether the pattern of TFBS disruption at the focal SNPs was specific to these variants or typical of *cpo* intronic SNPs more broadly, we compared TFBS disruption rates across three groups: the three focal clinal SNPs, 41 additional clinal SNPs from the same region (CQ < 10^-4^), and 30 non-clinal SNPs drawn from the same 30kb intronic window. Focal clinal SNPs did not differ from other *cpo* SNPs in overall number of disrupted motifs (focal mean 9.3 vs. other clinal 8.8 vs. non-clinal 10.0; Kruskal-Wallis p=0.783), indicating they are not generally more disruptive of transcription factor binding. However, focal SNPs were significantly enriched for *EcR::usp* and *cnc::maf-S* binding site disruption relative to all non-focal SNPs combined (*EcR::usp*: 2 of 3 focal vs. 6 of 71 non-focal, Fisher’s exact p=0.029; cnc::maf-S: 2 of 3 focal vs. 3 of 71 non-focal, Fisher’s exact p=0.011). Notably, all *EcR::usp* and *cnc::maf-S* hits occurred exclusively in clinal variants (0 of 30 non-clinal SNPs from the same intronic region disrupted either motif), which is consistent with ecdysone and NRF2 pathway regulation being specifically associated with climatic adaptation at this locus rather than background sequence variation.

Extending this comparison across all 105 testable JASPAR motifs among the 44 clinal SNPs (non-clinal SNPs lack a defined CQ value and were excluded from this analysis), defined as motifs with at least two disrupting and two non-disrupting SNPs among the 44 clinal SNPs, *cnc::maf-S* ranked among the most strongly associated with clinal significance (3rd of 105; Wilcoxon p=0.029). The two motifs ranking above it (*CG32105*, *onecut*) were each supported by the minimum testable sample size in this analysis of only two disrupting SNPs and are not interpreted as stronger competing signals given their comparably fragile support. *EcR::usp* disruption did not reach significance in this same motif-wide framework (21st of 105; Wilcoxon p=0.197) despite having the largest sample size among top-ranked motifs (n=8 disrupting SNPs), though the direction of effect was consistent (higher clinal significance among disrupting SNPs). This suggests the *EcR::usp* and *cnc::maf-S* enrichment at focal SNPs reflects specific sequence context rather than a general consequence of strong clinal selection. No curated cis-regulatory modules from the REDfly database (v9.6.1; Gallo et al. 2023) overlapped the focal SNP positions, indicating these represent novel predicted regulatory elements not previously characterized.

### Genotype frequency distributions in natural populations

The positive epistasis observed for temperate and tropical genotypes suggests these combinations may be under environment-specific selection, which should be reflected in their frequencies in natural populations. We examined wild populations across spatial and temporal gradients, comparing observed genotype frequencies to a null expectation based on individual allele frequencies at each locus and the assumption of random assortment among focal loci. The null expectation assumed random assortment among loci and did not account for pairwise linkage disequilibrium, which varied among populations (Table S7). Because linkage disequilibrium in wild *D. melanogaster* generally decays over distances shorter than the 1–2kb separating the focal loci (Langley et al. 2012; Mackay et al. 2012), elevated genotype frequencies above the null expectation are consistent with positive selection or epistatic fitness effects maintaining the intact combinations.

Genotype frequencies across the latitudinal cline were consistent with environment-specific selection. Temperate and tropical genotypes showed opposite frequency clines from Florida (25.5°N) to Maine (44.0°N), with the tropical *cpo^CGT^* more common at low latitudes and the temperate *cpo^TTA^* highest at the northernmost site (Figure 5B). Both genotypes occurred more frequently than predicted from individual allele frequencies, consistent with positive selection maintaining the intact combinations (Figure 5B, Table S8). Both *cpo*^CGT^ and *cpo*^TTA^ showed a significant decline in deviation from the null prediction with increasing latitude (*cpo*^CGT^: F(1,2)=336.3, p=0.003, R²=0.994; *cpo*^TTA^: F(1,2)=38.3, p=0.025, R²=0.950), with the largest excess above prediction for both genotypes occurring at the southernmost site.

**Figure 5.**
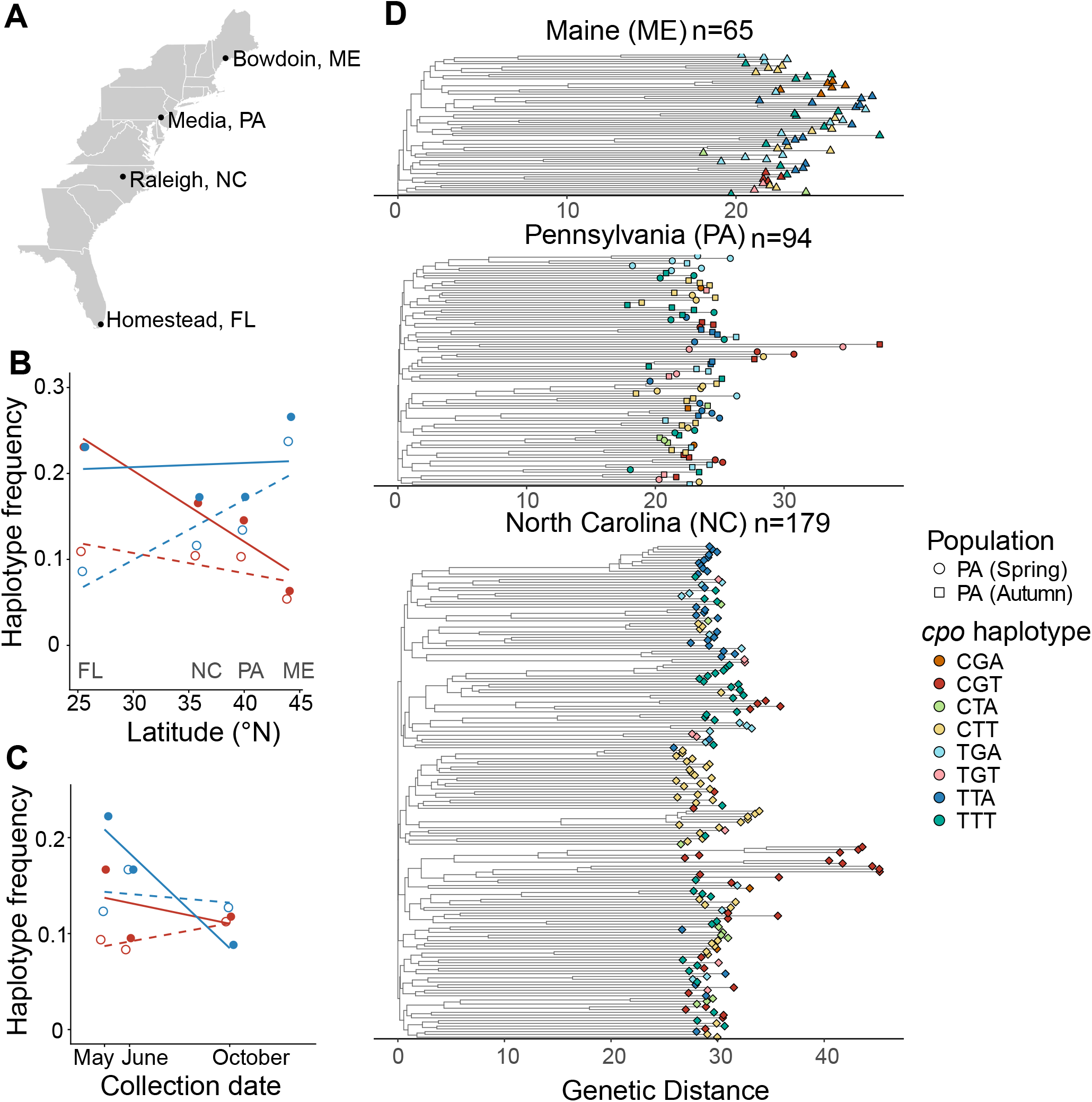
Geographic and temporal distribution of cpo haplotypes in wild *D. melanogaster* populations. (A) Collection sites (Homestead, FL to Bowdoin, ME). Temperate (*cpo*TTA, blue) and tropical (*cpo*CGT, red) haplotype frequency observed (solid/filled) compared to null expectation (dashed/open) across (B) latitude and (C) seasonal time in Pennsylvania. (D) Neighbor-joining trees of the ∼4kb focal region, by population; tips colored by haplotype, shaped by season (PA only).

Seasonal genotype frequencies in a Pennsylvania orchard similarly supported environment-specific selection (Figure 5C). As predicted from environmental parallels between seasonal and clinal gradients, temperate *cpo^TTA^* was more common than tropical *cpo^CGT^* in spring, and the pattern reversed in autumn. Temperate frequency was above the null prediction post-winter and declined below it in autumn, while the tropical genotype showed the complementary pattern; however, neither deviation was statistically significant over the five-month sampling period (temperate: F(1,2)=4.49, p=0.16; tropical: F(1,2)=0.61, p=0.52), likely reflecting the limited power of three seasonal time points.

### Haplotype distribution in the wild

The elevated frequencies of temperate and tropical haplotypes in natural populations could reflect either restricted recombination in the region surrounding the focal loci or selection acting on the genotype combinations. If recombination were restricted, SNPs located between the focal loci would be expected to be fixed in association with specific haplotypes. If instead differential selection shaped haplotype frequency, intervening SNPs would be free to recombine, producing heterogeneous backgrounds within each focal genotype class, unless selection itself locally maintains linkage. We used a 4kb haplotype spanning the focal loci and a permutation-based test of phylogenetic clustering to ask whether tips carrying each genotype combination sit closer together on a neighbor-joining tree than expected by chance, examining this separately within each population to avoid confounding with broader population structure (Figure 5D).

The tropical *cpo*^CGT^ haplotype showed no evidence of phylogenetic clustering in any population with sufficient sample size to test (Pennsylvania: n=12, permutation p=0.997; North Carolina: n=31, p=1.00), consistent with recombination continually regenerating this combination from a heterogeneous genomic background rather than descent from a single haplotype block. The temperate *cpo*^TTA^ haplotype showed a different, population-specific pattern: no significant clustering in Maine (n=13, p=0.39) or Pennsylvania (n=13, p=0.078), but significant clustering in North Carolina (n=32, p=0.0001), suggesting that in at least one population, selection may be acting quickly enough, or recombination arising rarely enough, to maintain local linkage around this combination. The rare *cpo*^CGA^ haplotype that was too infrequent in the DGRP to construct a recombinant outbred population showed significant clustering in Maine (permutation p=0.0002) despite its low frequency, though sample sizes this small (n<10 in every population) limit confidence in this result. Together, these findings indicate that recombination and selection likely act with different relative strengths on different haplotype backgrounds at this locus, rather than a single shared mechanism governing all combinations.

## Discussion

Non-additive interactions among three SNPs in *cpo* produce phenotypes that exceed additive predictions for multiple traits corresponding to expected selective pressures in their respective climatic environments. The temperate *cpo^TTA^* and tropical *cpo^CGT^* genotypes, or genetic variants linked to them, show stronger than predicted stress resistance and fecundity respectively. Clinal genotype frequencies and haplotype structure provide additional, though partial and population-specific, evidence consistent with selection acting on these combinations; seasonal frequency shifts were directionally consistent with this pattern but did not reach statistical significance. Overall, our findings provide an example of pleiotropic life history tradeoffs shaped by non-additive interactions among intragenic polymorphisms, and demonstrate that the functional unit of adaptation at *cpo* is not a single SNP but a multi-allele combination shaped by selection acting differently across genetic backgrounds.

### Epistatic effects are robust across genetic backgrounds

The non-additive phenotypes observed in ROPs were largely recapitulated in Pennsylvania inbred lines fixed for each of the eight *cpo* genotype combinations, providing independent validation in an isogenic background. Temperate *cpo^TTA^* inbred flies survived starvation significantly longer than the additive prediction and longer than tropical *cpo^CGT^*flies, mirroring the ROP results. Tropical *cpo^CGT^* flies recovered from chill faster than temperate flies, consistent with the ROP pattern. Effect sizes were smaller in the inbred lines than in the ROPs, which is expected given that the isogenic PA background may suppress or amplify epistatic effects relative to the heterogeneous DGRP-derived backgrounds used in the ROPs. The directional concordance across both experimental systems strengthens the inference that the epistatic effects reflect intrinsic properties of the *cpo* genotype combinations rather than background-dependent artifacts. This is particularly important given that epistasis in natural populations is often background-dependent (Chandler et al. 2014), and the ROP design, while powerful, confounds genotype effects with the specific inbred line backgrounds used to construct each population.

### Pleiotropic life history tradeoffs

The broad patterns of phenotypic differences between temperate and tropical *cpo* genotypes are consistent with the classic life history tradeoff between stress resistance and reproductive output that characterizes clinal variation in *D. melanogaster* (Chippindale et al. 1996; Paaby and Schmidt 2009). As with all common-garden approaches, the fitness-associated phenotypes measured here were assayed under stable, resource-abundant laboratory conditions and may not directly translate to fitness dynamics under the more heterogeneous and often more stressful conditions flies experience in nature. Temperate *cpo^TTA^*flies showed higher dormancy, starvation resistance, and lipid content, traits associated with overwintering survival, while tropical *cpo^CGT^* flies had higher fecundity, consistent with adaptation to environments where population growth rather than survival dominates selection. Pleiotropy contributes to the evolution of strong genetic correlations in life histories, including the negative associations produced by antagonistic pleiotropy when the same genetic element encodes both positive and negative fitness phenotypes (Williams 1957; Rose 1984). The selection regime imposed by cold temperate winters likely favors genotypes that facilitate survival, and other genetically correlated traits may evolve as coadapted responses or through indirect selection via pleiotropy (Paaby and Schmidt 2009).

One result that runs counter to simple climate-based predictions is the significantly shorter lifespan of temperate *cpo^TTA^*flies relative to tropical *cpo^CGT^* flies. This is surprising given that temperate flies are generally expected to show higher stress resistance and longevity. Notably, temperate flies also showed significant negative epistasis for lifespan and lived shorter than the additive prediction in addition to shorter than tropical flies, suggesting that the three-allele combination carries a specific cost to longevity beyond what individual alleles predict. This pattern may reflect a tradeoff between dormancy investment and longevity outside the dormancy context; the metabolic and physiological shifts that facilitate overwintering may come at a cost to lifespan under benign laboratory conditions. Alternatively, the lifespan difference may reflect antagonistic pleiotropy in which alleles that enhance stress resistance and dormancy reduce longevity through shared downstream pathways, consistent with the broader observation that dormancy and longevity do not always covary positively in *Drosophila* (Schmidt et al. 2005).

### Predicted regulatory mechanism: ecdysone and NRF2 pathways

SNPs segregating at *cpo*, or genetic variants linked to them, interact to regulate *cpo* expression and shape fitness-associated traits. Relative whole-body *cpo* mRNA expression in temperate females was approximately 3.8-fold lower than in tropical females, and while the deviation of temperate flies from the additive prediction did not reach significance (p=0.093), the large effect size (Cohen’s d=2.49) suggests this reflects limited statistical power rather than a true null.

The TFBS scan provides a molecular hypothesis connecting the focal SNPs to *cpo* regulation. The convergent creation *of EcR::usp* binding sites at both *cpo^5558^* and *cpo^7866^* each independently targeting position 7 of the motif suggests the temperate haplotype has been under selection to place *cpo* expression under ecdysone receptor control. *EcR::usp* functions as a heterodimer that can either activate or repress target genes depending on the presence of its ligand, 20-hydroxyecdysone (20E). In the absence of 20E, as under standard laboratory conditions, unliganded EcR acts as a transcriptional repressor by recruiting corepressors to its bound sites (Tsai et al. 1999; Ghbeish et al. 2001). This repressor activity would be consistent with the lower *cpo* expression observed in temperate ROPs under standard conditions. When 20E titers rise during dormancy preparation, the repressor complex would be displaced and replaced with an activator complex, potentially upregulating *cpo* specifically at the onset of reproductive dormancy, consistent with the tissue-specific and dormancy stage-specific upregulation of *cpo* documented in diapausing dipterans (Schmidt et al. 2008; Zhang and Denlinger 2011; Salminen et al. 2015). The *Broad* binding sites present at *cpo^5558^* and *cpo^7866^* in the temperate background reinforce this interpretation: *Broad* is a primary ecdysone-responsive transcription factor that mediates the transcriptional cascade downstream of ecdysone signaling (Bayer et al. 1996), and its co-occurrence alongside *EcR::usp* sites suggests the temperate haplotype recruits multiple elements of the ecdysone signaling network to regulate *cpo*.

The *cnc::maf-S* result points to a second potential regulatory pathway. *CncC*, the *Drosophila* ortholog of *NRF2*, activates the oxidative stress response and helps maintain proteostasis, and is a well-established regulator of stress tolerance and lifespan: knocking down *CncC* reduces both in *Drosophila* (Sykiotis and Bohmann 2008; Pomatto et al. 2017). The temperate haplotype retains (at *cpo^4408^*) or creates (at *cpo^5558^*) *cnc::maf-S* binding sites absent in the tropical haplotype, raising the possibility that *CncC* activity contributes to the higher starvation resistance and lipid content observed in temperate flies. The shorter lifespan of temperate flies is counterintuitive given their stress resistance, but may reflect a dosage effect: moderate *CncC* activation is beneficial, while constitutive or excessive activation is detrimental to lifespan (Pomatto et al. 2017).

Together, these results provide evidence that the temperate haplotype may be regulated through ecdysone signaling or the antioxidant stress response, two pathways relevant to dormancy and seasonal adaptation. Neither the *EcR::usp* nor the *cnc::maf-S* sites are present in the tropical haplotype or at non-clinal *cpo* SNPs; the regulatory module emerges specifically from the temperate allele combination, offering a plausible, though currently untested, molecular hypothesis for the higher-order epistasis documented phenotypically. Because RNA-binding proteins broadly regulate post-transcriptional processes from splicing to translation (Lukong et al. 2008), lower *cpo* expression under this model could have wide-ranging pleiotropic effects. This is consistent with the tissue- and stage-specific expression of *cpo* documented in *D. melanogaster* and other dipterans (Schmidt et al. 2008; Zhang and Denlinger 2011; Salminen et al. 2015; Zhao et al. 2015), suggesting the focal SNPs affect transcript abundance in specific tissues or developmental stages rather than uniformly.

Studies of epistasis in natural populations have largely focused on interactions between genes or among amino acid changes within proteins (Natarajan et al. 2013; Hanifin and Gilly 2015; Tufts et al. 2015; Liu et al. 2019; Taverner et al. 2019; Borne et al. 2026); our results extend this to non-coding regulatory SNPs within a single gene. Intergenic epistasis has also been shown to shape polygenic adaptation in *Drosophila* (Forsberg et al. 2025), suggesting the same combinatorial logic may operate at the intragenic level as well.

### Fitness landscape and the adaptive walk

Visualizing the deviation from additive predictions across all the *cpo* genotype combinations reveals that non-additive effects are concentrated at the temperate *cpo^TTA^* and tropical *cpo^CGT^*vertices of the genotypic landscape, with intermediate genotypes generally showing weaker or opposite-signed deviations (Figure 3); this pattern was clearest for fecundity and dormancy. This pattern is inconsistent with a stepwise adaptive walk model, in which successive allele substitutions along a path from tropical to temperate would each confer increasing fitness and produce deviations distributed across intermediate genotypes (Weinreich et al. 2005; Weinreich et al. 2006). Instead, the fitness benefits appear to emerge specifically from the intact three-allele combinations, suggesting that the combinations arose from standing genetic variation rather than sequential mutation and selection, although some exceptions suggest the underlying landscape may include additional local structure beyond a simple two-peak model. Empirical fitness landscapes across diverse systems show that epistasis commonly concentrates fitness effects at specific genotypic combinations rather than distributing them uniformly across intermediates, consistent with the pattern we observe here (Barrett and Schluter 2008; Bank 2022; Diaz-Colunga et al. 2023). In contrast to studies finding primarily additive architectures for some adaptive traits in *Drosophila* (da Silva Ribeiro et al. 2025), our results suggest that the genetic architecture of climatic adaptation at *cpo* is predominantly non-additive, at least for the focal SNP combinations examined here.

Adaptation from standing genetic variation is favored because beneficial alleles are immediately available, typically start at higher frequency than novel mutations, and have been pre-tested by previous selection (Hermisson and Pennings 2005; Barrett and Schluter 2008). Standing allelic variation may already be present across multiple genetic backgrounds, unlike a new mutation, which starts out confined to the single background in which it arose, increasing the opportunity for epistatic interactions to emerge (Barrett and Schluter 2008). The concentration of epistatic effects at the endpoints of the genotypic landscape is consistent with this interpretation: the temperate and tropical combinations represent fitness peaks accessible from standing variation, while intermediate genotypes represent neither peak.

An example of sign epistasis that illustrates the importance of genetic background is the dramatically different outcomes produced by the allele at *cpo*^7866^ depending on the other focal loci. When both other focal loci carry temperate alleles, the substitution of T for A at *cpo^7866^*changes the genotype from *cpo^TTT^* to the highly stress-tolerant temperate *cpo^TTA^*; but in the tropical background, the identical substitution modifies the highly fecund *cpo^CGT^*to the rare *cpo^CGA^*. The genetic background thus fundamentally alters the phenotypic consequence of the same allele substitution. Similar sign epistasis may explain why *cpo^7866^* is associated with reproductive dormancy in North American *D. melanogaster* but not in European or Australian populations (Schmidt et al. 2008; Lee et al. 2011; Zonato et al. 2016); the epistatic effect of *cpo^7866^* depends on the allelic state at the other focal loci, which may differ in frequency across continents.

### Selection maintains intact combinations despite recombination

The haplotype distribution in wild populations supports selection, not just restricted recombination, shaping the elevated frequencies of temperate and tropical genotypes. The pattern differs by genotype. The tropical *cpo*^CGT^ haplotype showed no evidence of phylogenetic clustering in any population tested, consistent with recombination continually regenerating this combination from a heterogeneous genomic background. The temperate *cpo*^TTA^ haplotype, by contrast, clustered significantly in North Carolina, and the rare *cpo*^CGA^ haplotype clustered significantly in Maine despite its low frequency. Both are more consistent with selection maintaining local linkage in some populations than with uniform free recombination across all genotypes, suggesting recombination and selection act with different relative strengths on different haplotype backgrounds, consistent with the low but non-random pairwise linkage disequilibrium among the focal SNPs.

The *cpo* alleles appear to shuffle such that any combination can occur in natural populations, but selection on the resulting phenotypes shapes the frequency of each combination, and in some cases may locally counteract the homogenizing effect of recombination. The *cpo* haplotype structure suggests that co-selection on epistatic interactions is not sufficiently strong to prevent recombination from breaking up allelic correlations everywhere (Lewontin 1974; Du et al. 2015; Knief et al. 2019), yet the fitness benefits of the intact combinations are strong enough to maintain their elevated frequency, and in specific populations, to produce detectable genealogical clustering.

We hypothesize that positive epistasis among the temperate and tropical genotypes elevates their frequency in respective environments, while negative epistasis purges *cpo*^CGA^ from populations (Chae et al. 2014; Sohail et al. 2017; Behrman et al. 2018; Kuzmin et al. 2018). The significant clustering of *cpo*^CGA^ in Maine may reflect very recent origin or a smaller number of independent recombination events than for other rare haplotypes. The selection for both temperate and tropical *cpo* genotypes across all sampled environments may reflect the alternative selection pressures imposed by environmental heterogeneity across temporal scales. Seasonal changes in the relative importance of fitness traits, combined with antagonistic pleiotropy, can result in segregation lift that maintains polymorphism at many loci (Wittmann et al. 2017; Bertram and Masel 2019a; Bertram and Masel 2019b) and may facilitate rapid climatic adaptation in *D. melanogaster*.

The seasonal frequency data provide an additional window into this combinatorial architecture. The temperate *cpo*^TTA^ haplotype is more common than predicted post-winter and declines in autumn, while the tropical *cpo*^CGT^ shows the complementary pattern, directionally consistent with alternative selection pressures across the seasonal cycle. Population genomic analyses of the same Pennsylvania population identify two independent temporal modes of seasonal selection: a concurrent mode in which allele frequencies respond to the current season’s environment, and a lagged mode in which one season’s environment predicts allele frequency change in the following season(Behrman and Schmidt 2022). These modes operate on largely non-overlapping genomic substrates, indicating that they represent genuinely distinct evolutionary processes rather than different views of the same signal. *cpo* appears in both temporal modes, suggesting its regulatory variants contribute to both immediate seasonal tracking and cross-season carry-over effects. The maintenance of both temperate and tropical haplotypes across all sampled environments may reflect the alternative selective pressures imposed by these two temporal modes simultaneously, with concurrent selection favoring stress-resistant temperate combinations post-winter and lagged effects carrying environmental information across seasons in ways that maintain diversity at both endpoints of the genotypic landscape.

### Broader implications

Our results have implications beyond *cpo* for how we interpret the large numbers of adaptive-candidate SNPs identified in population genomic studies. If SNPs within a gene interact epistatically despite low linkage disequilibrium, each variant may appear independently adaptive when examined in isolation, inflating the apparent number of causal loci when the true fitness effect belongs to the combination. The three focal SNPs studied here represent a small subset of the 45 highly clinal *cpo* variants, each of which could plausibly participate in additional epistatic combinations not captured by our three-locus design, raising the possibility that the large numbers of apparently independently adaptive loci identified across species reflect a smaller number of underlying combinatorial units (Bergland et al. 2014; Kapun et al. 2021; Bank 2022). Comprehensive functional investigation of individual candidate loci, of the kind undertaken here, is necessary to distinguish independent adaptive variants from components of epistatic combinations.

## Materials and methods

### Genomic Analysis

The three focal SNPs were among the most significantly clinal (CQ < 10^-4^) *cpo* SNPs across the east coast of North America (Bergland et al. 2014). The SNP associated with female reproductive dormancy (Schmidt et al. 2008), 3R:17967866 (*cpo^7866^*), was highly clinal (CQ < 2×10^-5^). *cpo*^7866^ was originally identified as a non-synonymous Lys/Ile change in the RH splicing variant of exon 5 (Bellen et al. 1992); however, alternative analysis from the modENCODE classification of functional elements does not identify this splice variant and reclassifies this nucleotide position as an intron 36 nucleotides downstream of the 3’ end of exon 5 (Celniker et al. 2009; Graveley et al. 2011). It therefore remains uncertain whether *cpo*^7866^ represents an intronic SNP or an amino acid change, but the SNP is strongly associated with reproductive dormancy and other life history traits in North American *D. melanogaster* (Schmidt et al. 2008). We examined *cpo*^7866^ in combination with the two most significantly clinal SNPs, 3R:17964408 (*cpo*^4408^) and 3R:17965558 (*cpo*^5558^), which are located in an intron upstream of *cpo*^7866^ and may regulate *cpo* expression to influence fitness traits. Linkage disequilibrium (LD) among the 44 most clinal SNPs (CQ < 0.0001) was calculated directly from genotype calls in the 205 sequenced inbred lines of the Drosophila Genetics Reference Panel (DGRP;(Mackay et al. 2012)) using variant calls from a *cpo*-region VCF subset read into R (*VariantAnnotation* Bioconductor package, (Obenchain et al. 2014)). Each SNP pair was treated as biallelic presence/absence of the alternate allele for the purpose of estimating D’. For each pair of SNPs with genotype calls in at least 10 DGRP lines, D was calculated as the difference between the observed joint frequency of both alternate alleles and the product of their marginal frequencies, and normalized to D’ by dividing by the appropriate D_max (the maximum possible value of D given the marginal allele frequencies (Lewontin 1964)). Pairwise D’ values were visualized as a rotated triangular heatmap.

### Recombinant outbred populations

We examined the fitness consequences of the three focal loci in combination by comparing flies carrying all three alleles at higher frequency in temperate environments (temperate genotype, *cpo*^TTA^) with flies carrying the alleles more common in tropical environments (tropical genotype, *cpo*^CGT^). The functional effects of each genotype were assessed using recombinant outbred populations (ROPs), each containing one of the eight possible three-allele combinations in a heterogeneous genomic background. Each ROP was initiated using nine sequenced DGRP inbred lines that shared one of the three-allele combinations at the focal SNPs (Table S2). Ten gravid females from each founder line were pooled and allowed to lay eggs for 48 hours, at which point the females were removed. ROPs were maintained at a population size of approximately 2000 adults for more than 10 non-overlapping generations on standard cornmeal-molasses food at 25°C and 12L:12D photoperiod. Each ROP remained fixed for its three-allele genotype combination along with any physically linked loci (Paaby et al. 2014; Behrman et al. 2018; Durmaz et al. 2019; Betancourt et al. 2021). The ROP design allows testing of compositional epistasis using combinatorial allele substitutions in a heterogeneous background, thereby avoiding the confounding effects of higher-order epistasis that can arise in isogenic backgrounds. ROPs were created for seven of the eight three-allele genotype combinations; the *cpo*^CGA^ combination was not created because it occurred too rarely in the DGRP to identify sufficient founder lines (Table S8).

### Phenotypic Assays

#### Life history traits

Fitness-associated life history traits were measured using flies from the ROPs. All experimental populations were reared at low density (∼30 eggs per vial) on standard cornmeal-molasses medium at 25°C, 12L:12D in Percival I36VL incubators. Traits were measured on individuals of both sexes at 3–5 days post-eclosion unless otherwise noted.

#### Reproductive dormancy

Approximately 50 females of each genotype were collected within 2 hours of eclosion and placed at 11°C, 10L:14D in Percival I36VL incubators for four weeks. Flies were then dissected to assess ovarian developmental status (King 1970); reproductive dormancy was defined by the absence of ovarian vitellogenin. Individuals with vitellogenic egg chambers were scored as non-dormant (Saunders et al. 1989; Williams and Sokolowski 1993; Tatar et al. 2001; Denlinger 2002; Schmidt and Paaby 2008). Although the appropriateness of the previtellogenic cutoff has been questioned because some flies may have vitellogenic egg chambers upon eclosion before shifting to dormancy conditions (Lirakis et al. 2018), the flies dissected here had normally developed ovaries with the expected progression of egg chamber development in each ovariole, confirming that the previtellogenic cutoff was appropriate.

#### Lipid content

Dry weight and fat content of flies 24 hours post-eclosion were determined following (Robinson et al. 2000). Flies were sorted into single-sex groups of 3 females and 4 males for weighing. Dry weight was measured after 24 hours at 60°C, and flies were subsequently reweighed after lipids were removed by 24 hours of light shaking in diethyl ether followed by 24 hours drying at room temperature. Lipid content was calculated as the difference between dry weight and lipid-free weight.

#### Starvation resistance

Ten replicates from each ROP were sorted into groups of 10 by sex using light CO₂ anesthesia and allowed to recover on standard cornmeal-molasses food for 24 hours, then transferred to vials containing cotton and 2 mL deionized water. Mortality was recorded at three timepoints throughout each day until all flies died.

#### Lifespan and fecundity

Flies were collected upon eclosion and sorted by sex under light CO₂ anesthesia. Ten replicate demography cages per genotype were each initiated with 50 males and 50 females from a single 24-hour cohort. Populations were maintained at 25°C, 12L:12D, and food plates were changed daily, at which point both eggs laid and adult mortality were scored to calculate average daily reproductive output per female.

#### Thermal tolerance

Response to high and low temperatures was measured for sixteen flies of each sex per genotype using methods previously described (Behrman et al. 2015). Flies were placed into individual glass tubes under light CO₂ anesthesia and given one hour to recover. Chill coma recovery time was measured by incubating the tubes at 4°C for two hours, then estimating recovery as the time required for each fly to resume an upright stance and locomotor activity, measured every 10 seconds using DAM2 activity monitors (TriKinetics, Waltham, MA) for one hour. Heat knockdown time was evaluated by placing vials at 25°C in a Percival I36VL incubator programmed to increase temperature by 1°C per minute until reaching 37°C, with knockdown estimated as the time at which locomotor activity ceased as measured by the DAM2 monitors.

#### *cpo* expression

Quantitative RT-PCR was used to assess whether cpo transcriptional expression differed among genotypes. Mated female flies aged 3–5 days, cultured on standard cornmeal-molasses food at 25°C with 12L:12D photoperiod, were flash-frozen at -80°C in pools of 15 individuals six hours after the onset of the light cycle. RNA was isolated using the Qiagen RNeasy Kit with tissue homogenization using the Qiagen QIAshredder. Quality and quantity of RNA were assessed on a NanoDrop ND-1000 spectrophotometer (ThermoScientific). cpo was reverse transcribed using Universal Probe Library probe #145 (forward: ACGGCACAAAAACCAGTTAAA, reverse: TTGGCTATTTTGACGAGTCAGTT; Roche Life Science cat# 04694317001) to synthesize an 87 nt cDNA amplicon spanning exons 4 and 5 and detecting nine of the twelve cpo transcripts (cpo-RN, cpo-RO, cpo-RP, cpo-RR, cpo-RS, cpo-RT, cpo-RU, cpo-RX, and cpo-RY). cDNA was quantified using TaqMan qRT-PCR (ThermoFisher Scientific). Expression levels were standardized relative to two endogenous control genes, Rpll140 and RpL32 (Life Technologies, Dm02134593_g1 and Dm02151827_g1, cat #4331182), and ΔCq was calculated as the difference between cpo expression and the mean expression of the two controls.

### Inbred line analysis

To validate the epistatic effects observed in ROPs in an isogenic background, we assessed chill coma recovery, starvation resistance, and heat knockdown in Pennsylvania inbred lines fixed for known genotypes at all three focal SNPs. Lines were derived from a mid-Atlantic population collected in Media, PA (39.12°N) and inbred for approximately 20 generations using a paired-sibling mating scheme (see DNA sequencing section below). Genotypes at the three focal SNPs were determined from whole-genome sequencing (see below). Of 129 genotyped lines, heterozygous lines and lines with ambiguous calls were excluded, leaving lines fixed for one of the eight three-allele combinations. The number of lines per genotype after filtering was: *cpo*^CGT^ n=14, *cpo*^TTA^ n=14, *cpo*^CTT^ n=27, *cpo*^TTT^ n=15, *cpo*^TGA^ n=14, *cpo*^CTA^ n=6, *cpo*^TGT^ n=3, *cpo*^CGA^ n=2.

Phenotypic assays on inbred lines followed the same protocols as for the ROPs above, with the following trait-specific notes. For starvation resistance, the survival time was recorded as hours to death; individual flies that died within 24 hours of the start of the assay were excluded as likely handling-stress mortalities (n=342 removed). For chill coma recovery, the standard 2-minute minimum cutoff was applied; the minimum observed recovery time in this dataset was 2.5 minutes, so no individuals were excluded on this basis.

Epistasis in inbred lines was assessed using linear mixed models with line as a random effect to account for multiple measurements per line: lmer(response ∼ *cpo*^4408^ + *cpo*^5558^ + *cpo*^7866^ + Sex + (1|Line)) for the additive model and lmer(response ∼ Combination + Sex + (1|Line)) for the observed model, fitted using the lme4 package (Bates et al. 2015) in R.

### DNA sequencing

The distribution of three-allele genotype combinations in natural populations was determined from a combination of publicly available sequence data from wild-derived inbred lines and wild-derived lines made isogenic at the third chromosome.

Temperate flies collected from Bowdoin, ME (44.01°N) and mid-Atlantic collections from Media, PA (39.12°N) in May, June, and October were used to establish isofemale lines (Behrman et al. 2018). Lines were inbred for approximately 20 generations using a paired-sibling mating scheme to produce 123 viable, fertile lines. Pools of 30–50 individuals per line were homogenized in lysis buffer with 1 mm beads, DNA was precipitated and resuspended in TE buffer, and libraries were prepared with a modified Nextera protocol (Baym et al. 2015). Samples were indexed with Illumina TruSeq Dual Index Sequencing Primer Kit (PE-121-1003) and pooled into three equimolar sets of approximately 70 samples each. Each pool was purified with Ampure XP, size-selected to 450–500 bp with a SizeSelect E-Gel, amplified with an additional 5 rounds of PCR, and sequenced on the HiSeq 3000. Sequences are deposited in SRA (BioProject PRJNA383555). Adapter sequences were trimmed (Trimmomatic v0.33), overlapping reads were merged (PEAR v0.9.6), and reads were mapped to the D. melanogaster reference genome (v5.39) using BWA (v0.7.9) with default parameters. PCR duplicates were removed with PicardTools (v1.12) and GATK’s Unified Genotyper (v3.4-46) was used for base quality score recalibration, indel realignment, and SNP discovery.

The mid-Atlantic collection from Raleigh, NC (35.5°N) is represented by the publicly available DGRP sequenced inbred lines (Mackay et al. 2012).

Tropical flies from Jacksonville, FL (30.32°N) and Homestead, FL (25.5°N) were made isogenic at the third chromosome using the TM3 balancer. Ethanol-preserved flies were rehydrated in dH₂O for 24 hours and homogenized in digestion solution using a mini bead beater (Biosphere Products Inc.). Genomic DNA was extracted using the DNeasy kit (Qiagen) and three sections of cpo were amplified by PCR using primers designed with Primer3 (v4.0.0): for cpo4408, forward ACTTCCGTTTCAGCATAGTT, reverse TCATTCAGTTGAGCCAATGA; for cpo5558, forward AAAGCCCTGTTTGTATAGCA, reverse ATAAACATAGTGGGCCAGTG; for cpo7866, forward AATGGAGAATCAGATGGCTC, reverse AACAATTTGCATGGACCTTG. Fragments of approximately 600 nt (*cpo^4408^*), 500 nt (*cpo^5558^*), and 400 nt (*cpo^7866^*) were amplified in 40 cycles at 45°C annealing temperature. Products were purified using Qiagen PCR purification kits and sequenced in the reverse direction at the University of Pennsylvania DNA Sequencing Center on an ABI 3100 sequencer (Applied Biosystems). Sequences were aligned using Geneious 7.1.5 (Biomatters).

### Haplotype structure

Haplotype distribution was assessed separately in each of the three sequenced populations (Maine, Pennsylvania, and North Carolina) using all variants within approximately 4kb spanning the three focal SNPs. For Maine and Pennsylvania, genotypes were drawn from the inbred line panel described above; for North Carolina, genotypes were drawn from the DGRP. Genotype calls were converted to a numeric dosage matrix, and pairwise genetic distances among individuals were calculated and used to construct a neighbor-joining tree separately for each population using the *ape* package (Paradis and Schliep 2019) in R, avoiding confounding between genotype-specific clustering and broader population structure. A combined, unrooted neighbor-joining tree across all three populations was additionally constructed for visual reference (Figure S4), using the subset of SNPs shared across populations after coordinate harmonization between genome assembly versions.

To formally test whether tips carrying a given three-locus haplotype clustered more closely on each population’s tree than expected by chance, we calculated the mean pairwise patristic (tree) distance among same-haplotype tips and compared this to a null distribution generated by permuting haplotype labels across tips 9,999 times, holding tree topology and group size fixed. A one-sided permutation p-value was calculated as the proportion of permuted mean distances less than or equal to the observed value. This test was performed independently for each of the eight three-locus haplotypes within each population with at least two individuals carrying that haplotype. To confirm that this permutation test did not disproportionately flag larger haplotype groups as significantly clustered due to sample size alone, we performed a null-calibration check: for a range of group sizes matching those observed in the real data, we randomly assigned tips to fake groups with no true underlying structure, applied the identical permutation test procedure, and calculated the resulting false-positive rate across 200 replicate simulations per group size. False-positive rates remained at or below the nominal 5% threshold across all tested group sizes and were not positively associated with group size, confirming the clustering test is not biased toward larger groups.

### Statistical analysis

All analyses were performed in R v4.6.0 (R Core Team), with Claude (Anthropic, Sonnet 5) assisting with debugging and reformatting analysis code and generating figure alt text for accessibility; all statistical analyses, results, and interpretations were performed and verified by the authors.

For the ROP data, separate generalized linear models were fitted for the additive effects of individual SNPs and the combined genotype effect:

Additive model: Trait ∼ *cpo^4408^* + *cpo^5558^* + *cpo^7866^* (+ Sex where applicable) Observed model: Trait ∼ Combination (+ Sex where applicable)

The significance of each individual SNP’s effect within the additive model was assessed using Type II tests (car package; (Fox and Weisberg 2019). To complement the genotype-specific deviation tests described below, we additionally tested for genotype combination effects beyond additive main effects overall using a likelihood-ratio test comparing the additive model to the full Combination model for each trait. For starvation resistance, lifespan, and development time, a random effect for bottle/vial nesting was included in both models to account for non-independence of individuals sampled from the same replicate vial. For reproductive dormancy, binomial family GLMs were used; all other traits used Gaussian linear models. Marginal means (emmeans) and standard errors for each allele at each SNP position were estimated from the additive model using the emmeans package (Lenth and Piaskowski 2026) and are reported (Table S3). Stepwise model selection using stepAIC (package MASS) and the likelihood ratio test identified the most parsimonious model for each trait.

Epistasis was assessed by testing whether observed trait values deviated significantly from an additive null prediction. The null prediction for each three-allele combination was the mean of the three marginal emmeans, one for each SNP position, derived from the additive model. For example, the null prediction for *cpo^TTA^* was the mean of the emmean for the T allele at *cpo^4408^*, the T allele at *cpo^5558^*, and the A allele at *cpo^7866^*(Figure S1). For binomial traits, a one-sample proportion test (Pearson’s chi-square with continuity correction) was used to test deviation between observed and predicted proportions. For continuous traits with a bottle/vial random effect in the observed model (starvation resistance, lifespan, and development time), individual values were first averaged to bottle-level means, and a Welch one-sample t-test was used to compare these replicate means against the additive prediction; for other continuous traits, individual-level values were compared directly against the additive prediction using the same test. Effect sizes were calculated as Cohen’s d (effsize package, (Torchiano 2016)). The same approach was applied to inbred line data, with line-level emmeans from the observed lmer model as the observed values.

### Genotype frequency analyses

The predicted genotype frequency in each population was calculated from individual allele frequencies within that population under the assumption of random assortment: predicted frequency = frequency(allele 1) × frequency(allele 2) × frequency(allele 3). Standard errors for observed and predicted genotype frequencies were calculated as √(P(1−P)/n). The significance of deviations between observed and predicted genotype frequencies across the latitudinal cline was assessed by regressing the deviation (observed minus predicted) directly on latitude, separately for each genotype (Deviation ∼ Latitude). Seasonal deviations were assessed with the same model structure using collection date as the predictor.

### Transcription factor binding site analysis

To identify transcription factor binding sites (TFBS) predicted to be differentially affected by temperate and tropical alleles at the three focal SNPs, we extracted the genomic sequence in a ±20bp window centered on each focal SNP position (3R:17964408, 3R:17965558, 3R:17967866) from the *D. melanogaster* reference genome (dm6) using the BSgenome.Dmelanogaster.UCSC.dm6 Bioconductor package (v1.4.3). For each SNP we constructed two 41bp sequences differing only at the central position, one carrying the reference allele and one carrying the alternate allele. All 143 *D. melanogaster* transcription factor position weight matrices in the JASPAR2020 database (Fornes et al. 2020) were retrieved using TFBSTools (v1.36; (Tan and Lenhard 2016). Each PWM was scored against both allele sequences on both strands using the searchSeq function with a minimum relative score threshold of 80%, and the maximum relative score in the window was recorded for each allele. Score differences (ALT minus REF relative score) exceeding an absolute value of 0.1 were retained as putative TFBS disruptions.

For each retained SNP-motif pair, the hit coordinates and strand of the higher-scoring allele were used as a shared anchor to extract the corresponding sequence from both alleles, ensuring both alleles’ displayed binding-site sequences reflect an identical genomic window and orientation. For *EcR::usp* specifically, the position within the 15bp motif at which each SNP altered binding was determined from this shared anchor. The probability of two independent SNPs both targeting the same specific motif position by chance was calculated as (1/L)^2^, where L=15 is the motif length, giving p=0.0044.

To assess whether the pattern of TFBS disruption at the focal SNPs was specific to these variants or typical of *cpo* intronic SNPs more broadly, we compared TFBS disruption rates across three groups: the three focal clinal SNPs, 41 additional clinal SNPs from the same region (CQ < 10^-4^) not among the three focal loci, and 30 non-clinal SNPs drawn from the same intronic region (3R:17950000–17980000) present in the DGRP *cpo* region VCF but absent from the list of significantly clinal variants. The identical scanning procedure described above was applied to all SNPs in each group. A Kruskal-Wallis test was used to compare the overall number of disrupted motifs per SNP across the three groups. Fisher’s exact tests (one-tailed, greater) were used to test whether focal clinal SNPs were enriched for predicted *EcR::usp* and *cnc::maf-S* binding site disruptions relative to all non-focal SNPs (other clinal and non-clinal combined). An identical Fisher’s exact test was used to test enrichment for *Broad* complex binding site disruption, pooling across all annotated Broad isoforms.

To evaluate whether the EcR::usp and cnc::maf-S enrichment reflected these motifs specifically, rather than a general property of strongly clinal SNPs, we extended this comparison across all JASPAR2020 *Drosophila* motifs among the 44 clinal SNPs with a defined CQ value (focal plus other-clinal; non-clinal SNPs were excluded from this analysis as they lack a CQ value). Motifs were retained for this analysis if at least two clinal SNPs disrupted the motif and at least two did not, yielding 105 testable motifs. For each testable motif, a Wilcoxon rank-sum test compared clinal significance (−log10 CQ) between disrupting and non-disrupting SNPs, and motifs were ranked by the resulting p-value.

All TFBS analyses were performed in R v4.6.0. The REDfly database (v9.6.1; (Gallo et al. 2011) was queried for annotated cis-regulatory modules overlapping the focal SNP positions in dm6 coordinates; none were found.

## Author contributions

ELB, AOB, DAP and PS designed the experiment. ELB performed the experiment, did the analyses and wrote the manuscript. PS, AOB and Dap provided input on the manuscript.

## Acknowledgements

We would like to thank Annalise Paaby, Spencer Koury, and Walt Eanes for providing third-chromosome extracted samples from natural populations for sequencing. We would like to thank Camilo Khatchikian, Stephanie Seifert and Dustin Brisson for assistance and equipment during the sequencing, along with Aaron Leichty and Scott Poethig for assistance and equipment for the qRT-PCR. This work would not have been possible without the community supportive resources available at FlyBase (Thurmond et al. 2019) and for permission to collect Drosophila from Linvilla Orchards. This work was supported by National Science Foundation Graduate Research Fellowship DGE-0822 to ELB; National Institutes of Health R01GM100366 to PS; and National Institutes of Health R35GM118165 to DAP.

---

**Table S1.** *cpo* SNPs that are clinal with a q-value (CQ) of <10^-4^ and their allele frequency at sampled populations along a latitudinal cline. Focal alleles are highlighted in bold.

| Position<br>on 3R | CQ | Allele frequency at latitude (°N) |  |  |  |  |  |  |  |  |
| --- | --- | --- | --- | --- | --- | --- | --- | --- | --- | --- |
|  |  | 25.47 | 25.47 | 30.32 | 33.39 | 35.77 | 40 | 40 | 44.03 | 44.03 |
| 17939248 | $6.90 \times 10^{-05}$ | 0.48 | 0.36 | 0.32 | 0.14 | 0.10 | 0.04 | 0.16 | 0.05 | 0.04 |
| 17940255 | $3.00 \times 10^{-06}$ | 0.19 | 0.09 | 0.23 | 0.26 | 0.47 | 0.37 | 0.35 | 0.59 | 0.58 |
| 17940301 | $3.40 \times 10^{-05}$ | 0.25 | 0.14 | 0.21 | 0.25 | 0.34 | 0.38 | 0.30 | 0.62 | 0.40 |
| 17940310 | $4.80 \times 10^{-05}$ | 0.44 | 0.53 | 0.49 | 0.43 | 0.39 | 0.10 | 0.13 | 0.07 | 0.27 |
| 17943485 | $3.00 \times 10^{-06}$ | 0.51 | 0.62 | 0.42 | 0.36 | 0.19 | 0.14 | 0.08 | 0.09 | 0.13 |
| 17947180 | $9.00 \times 10^{-06}$ | 0.67 | 0.68 | 0.73 | 0.51 | 0.42 | 0.39 | 0.39 | 0.23 | 0.22 |
| 17955967 | $7.90 \times 10^{-05}$ | 0.44 | 0.67 | 0.44 | 0.20 | 0.17 | 0.18 | 0.16 | 0.12 | 0.10 |
| 17955974 | $4.20 \times 10^{-05}$ | 0.37 | 0.52 | 0.43 | 0.14 | 0.06 | 0.08 | 0.12 | 0.04 | 0.03 |
| 17956579 | $1.90 \times 10^{-05}$ | 0.47 | 0.43 | 0.40 | 0.32 | 0.20 | 0.13 | 0.21 | 0.07 | 0.10 |
| 17956683 | $5.00 \times 10^{-06}$ | 0.47 | 0.46 | 0.55 | 0.28 | 0.11 | 0.14 | 0.16 | 0.04 | 0.10 |
| 17956684 | $1.00 \times 10^{-06}$ | 0.46 | 0.48 | 0.54 | 0.27 | 0.10 | 0.13 | 0.14 | 0.02 | 0.10 |
| 17956708 | $1.00 \times 10^{-06}$ | 0.49 | 0.53 | 0.54 | 0.32 | 0.15 | 0.14 | 0.29 | 0.06 | 0.13 |
| 17956748 | $3.10 \times 10^{-05}$ | 0.54 | 0.48 | 0.51 | 0.33 | 0.16 | 0.12 | 0.33 | 0.13 | 0.19 |
| 17956840 | $4.70 \times 10^{-05}$ | 0.51 | 0.67 | 0.47 | 0.28 | 0.24 | 0.13 | 0.21 | 0.10 | 0.41 |
| 17956850 | $4.90 \times 10^{-05}$ | 0.49 | 0.62 | 0.45 | 0.26 | 0.27 | 0.13 | 0.18 | 0.09 | 0.41 |
| 17957171 | $3.30 \times 10^{-05}$ | 0.60 | 0.65 | 0.76 | 0.53 | 0.27 | 0.32 | 0.43 | 0.30 | 0.19 |
| 17957175 | $3.60 \times 10^{-05}$ | 0.60 | 0.67 | 0.76 | 0.50 | 0.26 | 0.30 | 0.41 | 0.31 | 0.21 |
| 17957180 | 4.90 x10<sup>-05</sup> | 0.62 | 0.67 | 0.74 | 0.50 | 0.32 | 0.30 | 0.41 | 0.31 | 0.19 |
| 17957310 | 6.00 x10<sup>-05</sup> | 0.59 | 0.53 | 0.45 | 0.40 | 0.39 | 0.12 | 0.13 | 0.15 | 0.13 |
| 17958784 | 4.70 x10<sup>-05</sup> | 0.44 | 0.47 | 0.34 | 0.11 | 0.10 | 0.06 | 0.13 | 0.05 | 0.17 |
| 17958792 | 5.50 x10<sup>-05</sup> | 0.46 | 0.52 | 0.39 | 0.13 | 0.13 | 0.09 | 0.22 | 0.07 | 0.23 |
| 17959043 | 6.00 x10<sup>-06</sup> | 0.47 | 0.56 | 0.38 | 0.14 | 0.13 | 0.03 | 0.16 | 0.05 | 0.07 |
| 17963438 | 2.00 x10<sup>-05</sup> | 0.76 | 0.91 | 0.84 | 0.66 | 0.70 | 0.84 | 0.73 | 0.43 | 0.50 |
| 17963448 | 2.00 x10<sup>-06</sup> | 0.82 | 0.93 | 0.86 | 0.68 | 0.67 | 0.84 | 0.73 | 0.43 | 0.47 |
| 17963734 | 7.40 x10<sup>-05</sup> | 0.43 | 0.38 | 0.34 | 0.19 | 0.18 | 0.08 | 0.12 | 0.06 | 0.04 |
| 17964408 | <1 x10<sup>-07</sup> | **0.82** | **0.75** | **0.57** | **0.59** | **0.51** | **0.54** | **0.35** | **0.13** | **0.56** |
| 17965538 | 6.10 x10<sup>-05</sup> | 0.41 | 0.26 | 0.22 | 0.37 | 0.56 | 0.55 | 0.46 | 0.68 | 0.61 |
| 17965558 | <1 x10<sup>-07</sup> | **0.20** | **0.32** | **0.43** | **0.47** | **0.63** | **0.71** | **0.67** | **0.81** | **0.80** |
| 17965930 | 2.50 x10<sup>-05</sup> | 0.31 | 0.43 | 0.44 | 0.72 | 0.59 | 0.70 | 0.73 | 0.79 | 0.71 |
| 17965958 | 1.40 x10<sup>-05</sup> | 0.21 | 0.49 | 0.50 | 0.80 | 0.63 | 0.70 | 0.70 | 0.83 | 0.75 |
| 17967866 | **2.00** x10<sup>-06</sup> | **0.23** | **0.03** | **0.12** | **0.27** | **0.50** | **0.41** | **0.42** | **0.61** | **0.52** |
| 17970970 | 1.10 x10<sup>-05</sup> | 0.65 | 0.71 | 0.70 | 0.66 | 0.54 | 0.35 | 0.56 | 0.25 | 0.35 |
| 17984073 | 3.30 x10<sup>-05</sup> | 0.13 | 0.21 | 0.40 | 0.47 | 0.45 | 0.45 | 0.40 | 0.68 | 0.75 |
| 17984172 | 7.00 x10<sup>-06</sup> | 0.22 | 0.10 | 0.35 | 0.68 | 0.50 | 0.36 | 0.43 | 0.73 | 0.68 |
| 17986523 | 9.10 x10<sup>-05</sup> | 0.01 | 0.08 | 0.14 | 0.17 | 0.37 | 0.40 | 0.26 | 0.36 | 0.47 |
| 17989198 | 5.30 x10<sup>-05</sup> | 0.62 | 0.56 | 0.58 | 0.41 | 0.24 | 0.30 | 0.33 | 0.24 | 0.19 |
| 17994067 | 5.70 x10<sup>-05</sup> | 0.08 | 0.05 | 0.19 | 0.25 | 0.30 | 0.23 | 0.35 | 0.44 | 0.44 |
| 17997305 | $4.00 \times 10^{-06}$ | 0.85 | 0.76 | 0.61 | 0.46 | 0.35 | 0.59 | 0.47 | 0.29 | 0.29 |
| 17998827 | $8.00 \times 10^{-06}$ | 0.61 | 0.50 | 0.42 | 0.37 | 0.12 | 0.26 | 0.21 | 0.07 | 0.24 |
| 18000687 | $4.60 \times 10^{-05}$ | 0.72 | 0.91 | 0.75 | 0.61 | 0.42 | 0.52 | 0.60 | 0.28 | 0.71 |
| 18001545 | $7.30 \times 10^{-05}$ | 0.43 | 0.47 | 0.43 | 0.25 | 0.21 | 0.13 | 0.08 | 0.05 | 0.11 |
| 18013015 | $7.00 \times 10^{-06}$ | 0.67 | 0.62 | 0.62 | 0.59 | 0.18 | 0.50 | 0.62 | 0.14 | 0.21 |
| 18013084 | $<1 \times 10^{-07}$ | 0.59 | 0.58 | 0.41 | 0.39 | 0.16 | 0.31 | 0.35 | 0.07 | 0.08 |
| 18013126 | $<1 \times 10^{-07}$ | 0.59 | 0.64 | 0.45 | 0.36 | 0.09 | 0.34 | 0.34 | 0.11 | 0.10 |
| 18019905 | $2.60 \times 10^{-05}$ | 0.89 | 0.75 | 0.72 | 0.50 | 0.57 | 0.28 | 0.47 | 0.36 | 0.19 |

**Table S2.**
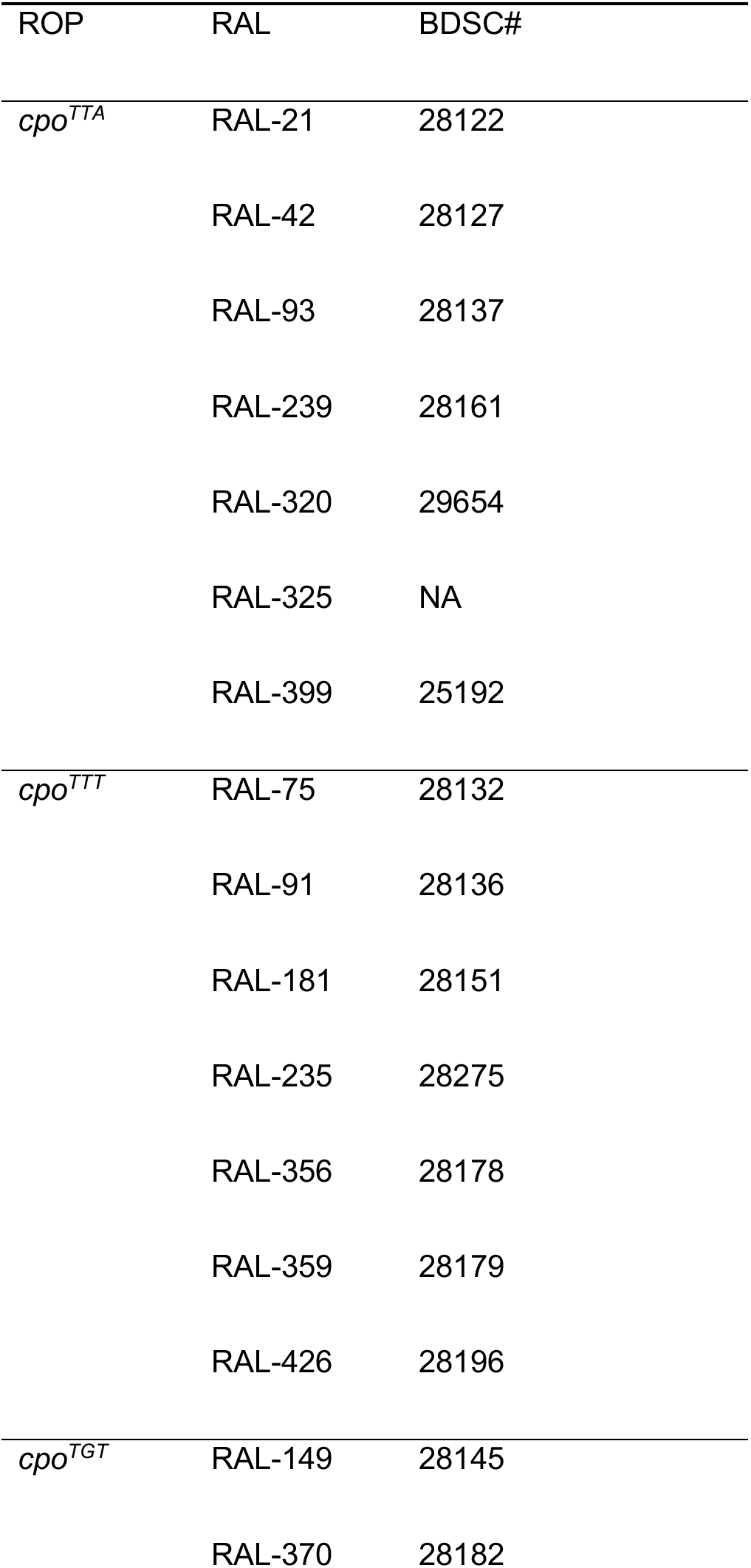
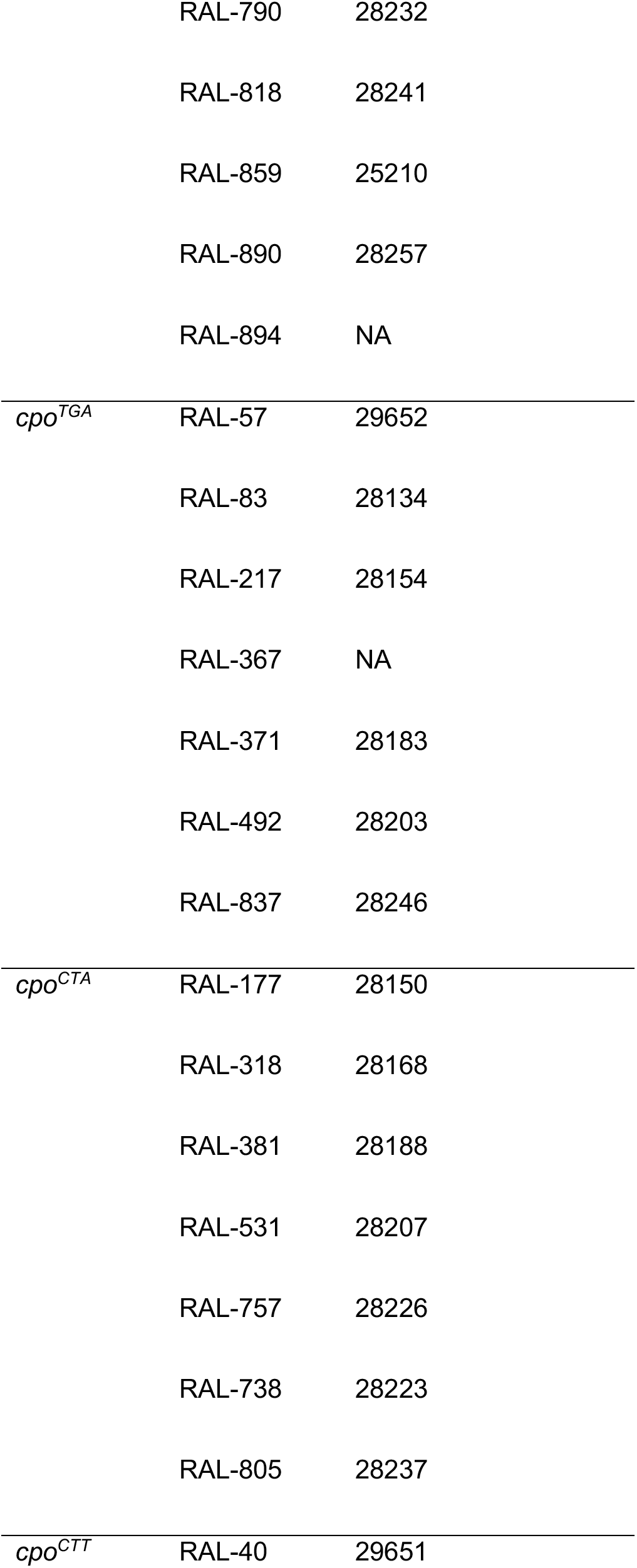
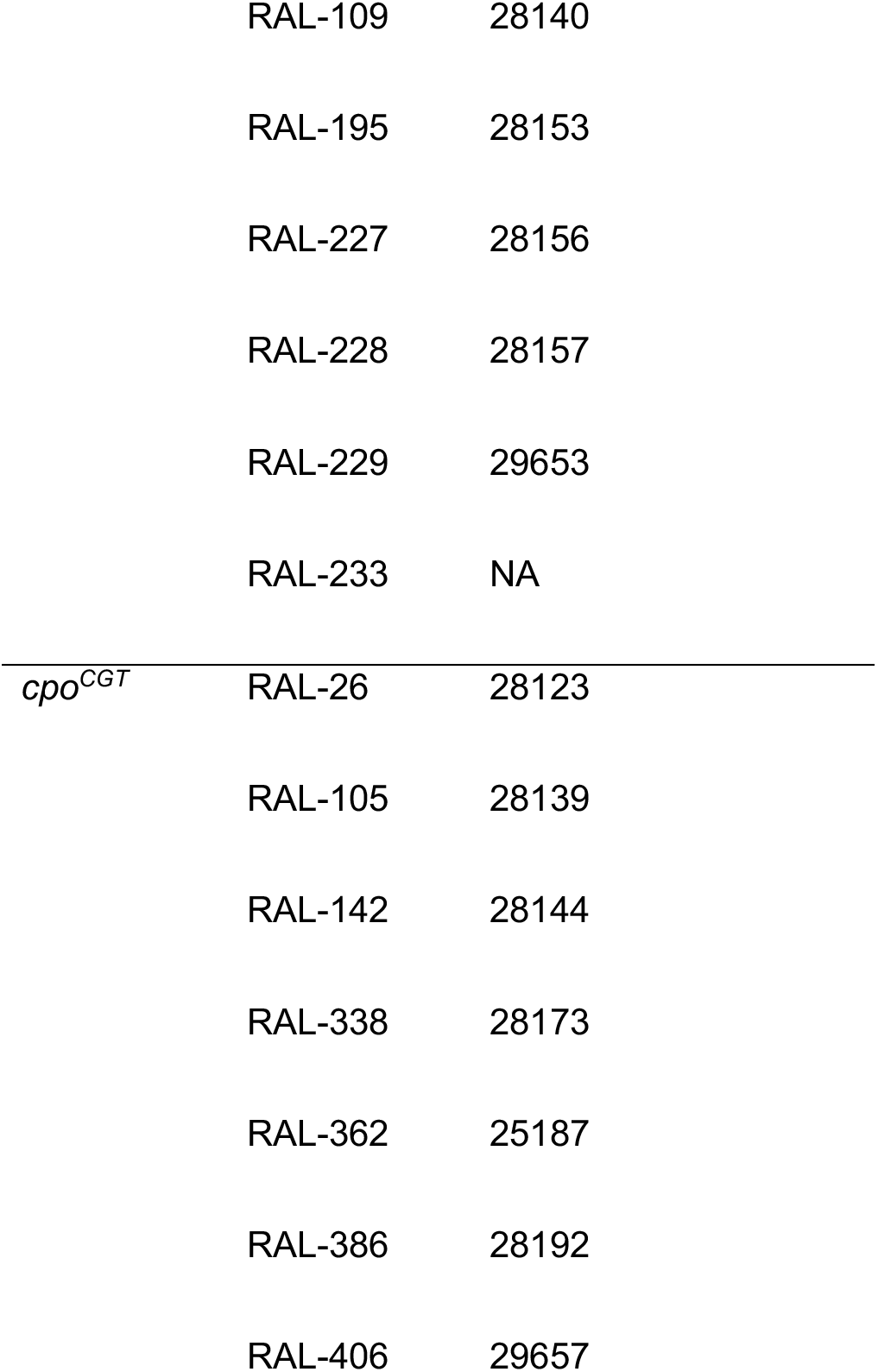
Drosophila Genetics Reference Panel lines (RAL) used to initiate each Recombinant Outbred Population. Bloomington Drosophila Stock Center number is indicated when available.

**Table S3.**
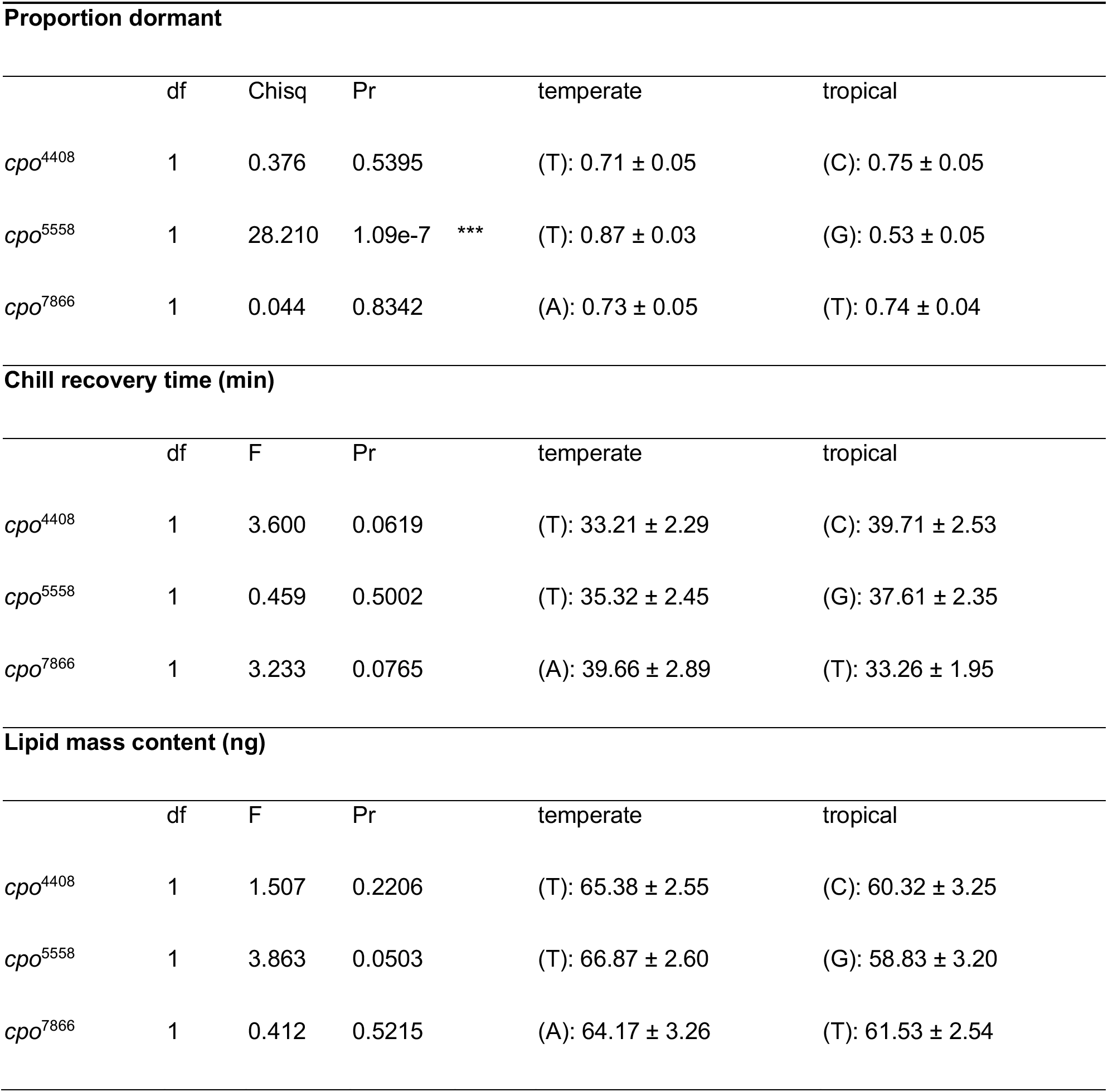
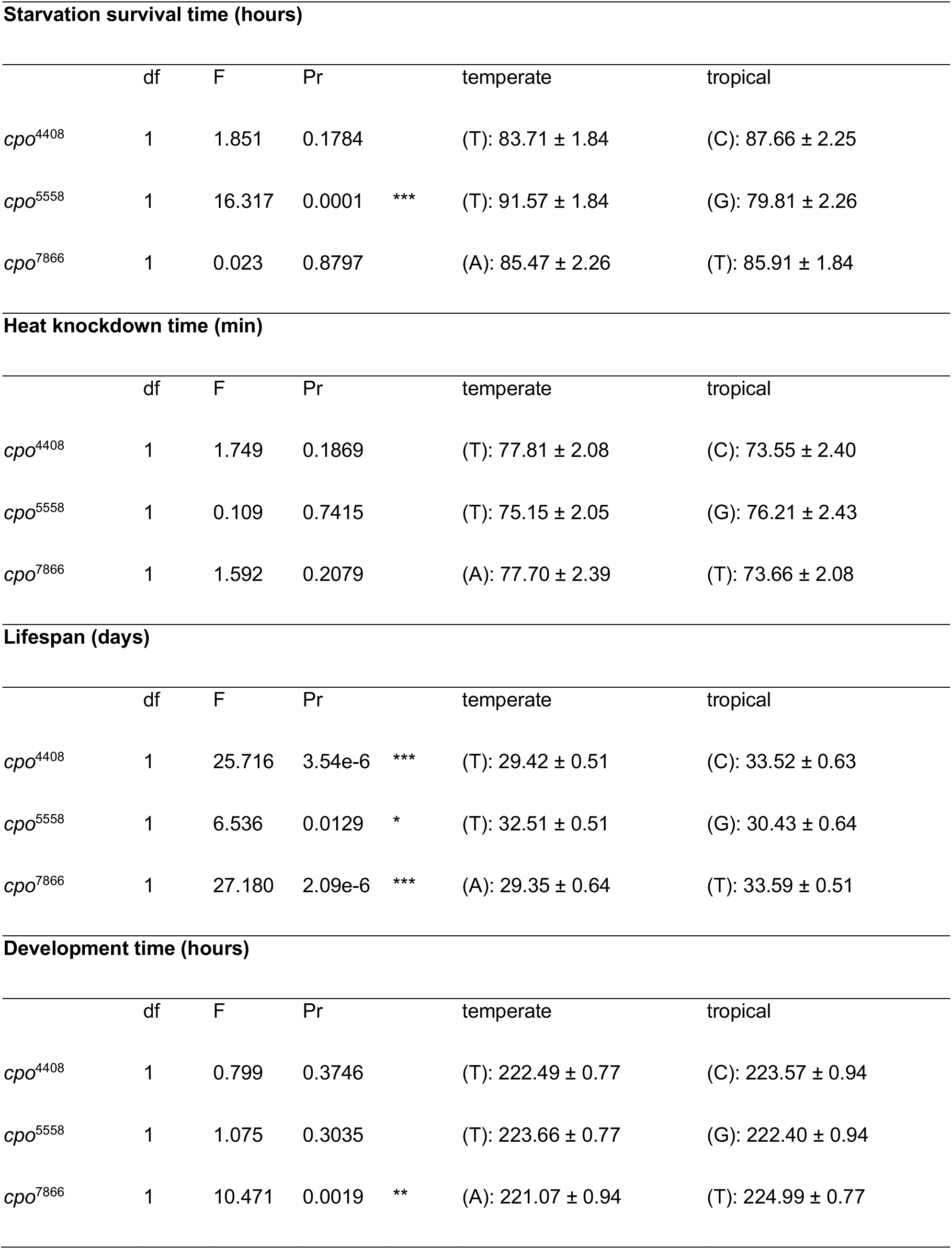
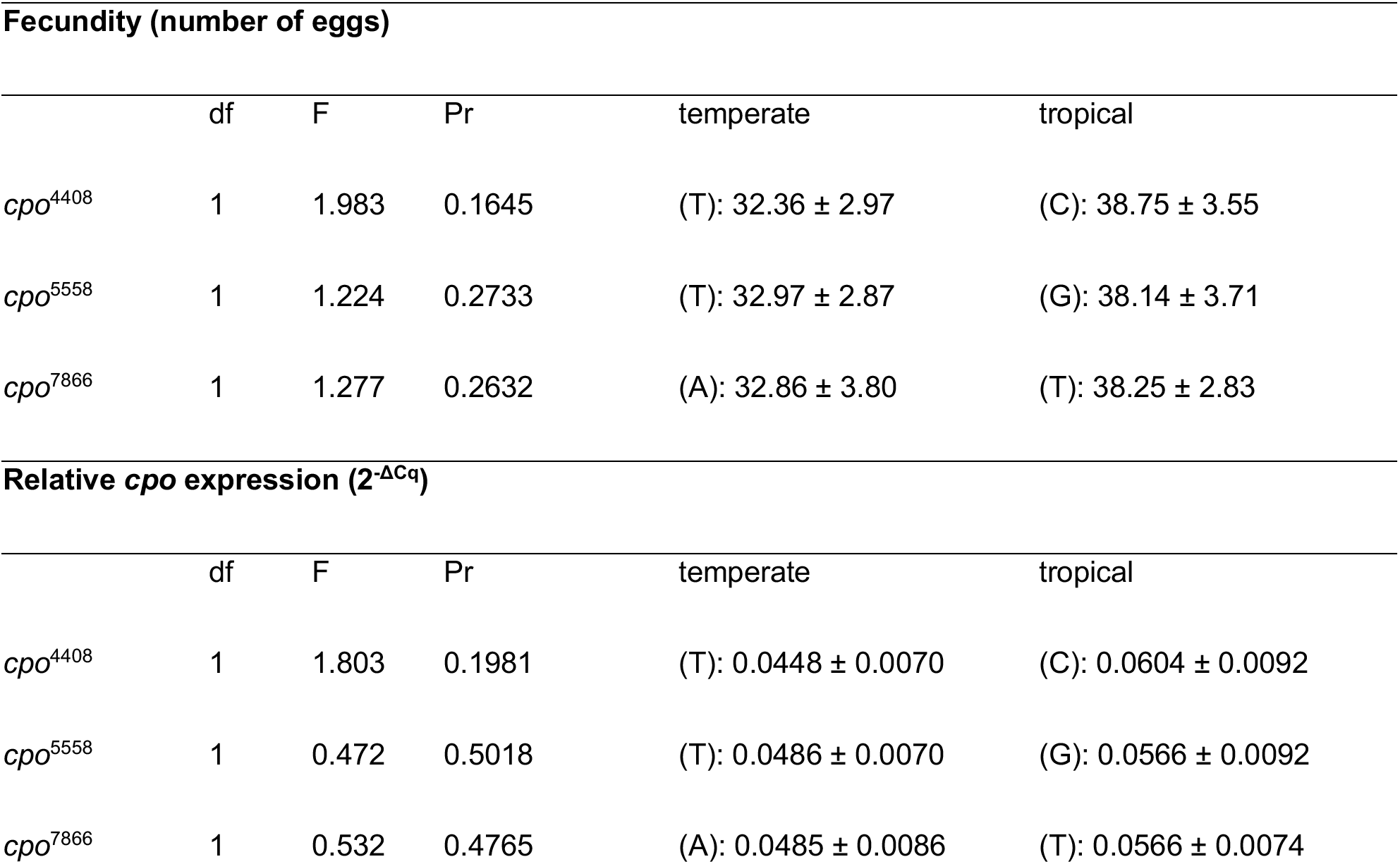
Effects of individual focal SNPs (*cpo*^4408^, *cpo*^5558^, *cpo*^7866^) on each fitness-related trait, from the additive model used to generate the null prediction for each three-locus genotype combination (see Methods). For each locus, the significance test is reported alongside the marginal mean ± SE for the temperate and tropical allele at that locus, estimated from the same fitted model. * p<0.05, ** p<0.01, *** p<0.001.

**Table S4.** Additive null prediction and observed trait value for each cpo three-locus genotype combination in recombinant outbred populations (ROPs). Prediction is the mean of the three marginal single-locus emmeans ± propagated SE; Observation is the ROP emmean ± SE. Deviation is observed minus predicted. The *cpo*^CGA^ combination is not shown, as it was too rare in the DGRP to establish a ROP.

| **Proportion dormant** |  |  |  |
| --- | --- | --- | --- |
| *cpo* haplotype | Prediction ± SE | Observation ± SE | Deviation |
| CGT | 0.674 ± 0.028 | 0.605 ± 0.079 | -0.069 |
| CTA | 0.784 ± 0.025 | 0.867 ± 0.051 | 0.082 |
| CTT | 0.789 ± 0.023 | 0.864 ± 0.052 | 0.075 |
| TGA | 0.655 ± 0.030 | 0.400 ± 0.089 | -0.255 |
| TGT | 0.660 ± 0.028 | 0.537 ± 0.077 | -0.124 |
| TTT | 0.775 ± 0.024 | 0.765 ± 0.102 | -0.011 |
| TTA | 0.770 ± 0.026 | 1.000 ± 0.500 | 0.230 |
| **Chill recovery time (min)** |  |  |  |
| *cpo* haplotype | Prediction ± SE | Observation ± SE | Deviation |
| CGT | 36.88 ± 1.89 | 41.81 ± 3.45 | 4.93 |
| CTA | 38.14 ± 1.48 | 35.73 ± 5.04 | -2.41 |
| CTT | 36.05 ± 1.90 | 33.93 ± 3.54 | -2.12 |
| TGA | 36.79 ± 1.90 | 36.65 ± 4.01 | -0.14 |
| TGT | 34.70 ± 1.48 | 28.31 ± 3.31 | -6.39 |
| TTT | 33.88 ± 0.65 | 30.25 ± 5.42 | -3.62 |
| TTA | 35.97 ± 1.89 | 44.68 ± 5.99 | 8.71 |

**Lipid mass content (ng)**
| *cpo* haplotype | Prediction ± SE | Observation ± SE | Deviation |
| --- | --- | --- | --- |
| CGT | 58.40 ± 0.96 | 55.79 ± 4.26 | -2.61 |
| CTA | 61.02 ± 1.80 | 60.67 ± 4.45 | -0.35 |
| CTT | 60.79 ± 1.82 | 60.54 ± 4.06 | -0.25 |
| TGA | 60.40 ± 1.82 | 57.20 ± 4.26 | -3.20 |
| TGT | 60.17 ± 1.80 | 60.17 ± 3.93 | -0.00 |
| TTT | 62.56 ± 1.19 | 64.49 ± 4.16 | 1.93 |
| TTA | 62.79 ± 0.96 | 70.78 ± 4.11 | 7.99 |

**Starvation survival time (hours)**
| *cpo* haplotype | Prediction ± SE | Observation ± SE | Deviation |
| --- | --- | --- | --- |
| CGT | 84.68 ± 2.34 | 86.86 ± 1.44 | 2.17 |
| CTA | 87.85 ± 1.81 | 82.10 ± 1.44 | -5.75 |
| CTT | 88.35 ± 1.41 | 100.51 ± 1.47 | 12.16 |
| TGA | 82.79 ± 1.41 | 70.83 ± 1.49 | -11.96 |
| TGT | 83.29 ± 1.81 | 81.52 ± 1.46 | -1.77 |
| TTT | 86.95 ± 2.21 | 76.57 ± 1.44 | -10.39 |
| TTA | 86.45 ± 2.34 | 105.35 ± 1.43 | 18.89 |

**Heat knockdown time (min)**
| *cpo* haplotype | Prediction ± SE | Observation ± SE | Deviation |
| --- | --- | --- | --- |
| CGT | 74.45 ± 0.92 | 70.97 ± 3.87 | -3.48 |
| CTA | 75.13 ± 1.14 | 73.61 ± 3.70 | -1.52 |
| CTT | 73.89 ± 0.37 | 73.22 ± 3.91 | -0.67 |
| TGA | 77.02 ± 0.37 | 84.40 ± 3.76 | 7.38 |
| TGT | 75.78 ± 1.14 | 73.10 ± 4.06 | -2.68 |
| TTT | 75.23 ± 1.15 | 77.51 ± 4.44 | 2.29 |
| TTA | 76.46 ± 0.92 | 74.65 ± 4.02 | -1.81 |

**Lifespan (days)**
| *cpo* haplotype | Prediction ± SE | Observation ± SE | Deviation |
| --- | --- | --- | --- |
| CGT | 32.55 ± 1.05 | 34.26 ± 1.02 | 1.71 |
| CTA | 31.84 ± 1.26 | 33.80 ± 1.00 | 1.97 |
| CTT | 33.25 ± 0.34 | 35.76 ± 1.00 | 2.51 |
| TGA | 29.77 ± 0.34 | 25.87 ± 1.06 | -3.90 |
| TGT | 31.18 ± 1.26 | 31.23 ± 1.00 | 0.05 |
| TTT | 31.88 ± 1.25 | 33.21 ± 1.00 | 1.33 |
| TTA | 30.47 ± 1.05 | 27.46 ± 1.00 | -3.01 |

**Development time (hours)**
| *cpo* haplotype | Prediction ± SE | Observation ± SE | Deviation |
| --- | --- | --- | --- |
| CGT | 223.67 ± 0.72 | 223.97 ± 1.31 | 0.31 |
| CTA | 222.84 ± 0.82 | 223.58 ± 1.31 | 0.73 |
| CTT | 224.09 ± 0.42 | 225.91 ± 1.30 | 1.83 |
| TGA | 222.03 ± 0.42 | 216.74 ± 1.31 | -5.29 |
| TGT | 223.28 ± 0.82 | 227.86 ± 1.30 | 4.58 |
| TTT | 223.70 ± 0.71 | 221.92 ± 1.32 | -1.78 |
| TTA | 222.45 ± 0.72 | 223.34 ± 1.31 | 0.88 |

**Fecundity (number of eggs)**
| *cpo* haplotype | Prediction ± SE | Observation ± SE | Deviation |
| --- | --- | --- | --- |
| CGT | 38.38 ± 0.19 | 53.26 ± 5.02 | 14.88 |
| CTA | 34.86 ± 1.95 | 25.69 ± 5.61 | -9.17 |
| CTT | 36.66 ± 1.85 | 35.52 ± 5.29 | -1.14 |
| TGA | 34.45 ± 1.85 | 24.44 ± 6.48 | -10.01 |
| TGT | 36.25 ± 1.95 | 32.59 ± 5.29 | -3.66 |
| TTT | 34.53 ± 1.87 | 30.60 ± 5.29 | -3.93 |
| TTA | 32.73 ± 0.19 | 37.99 ± 5.02 | 5.26 |

**Relative *cpo* expression ( $2^{-\Delta C_q}$ )**
| *cpo* haplotype | Prediction ± SE | Observation ± SE | Deviation |
| --- | --- | --- | --- |
| CGT | 0.0578 ± 0.0013 | 0.0589 ± 0.0157 | 0.0010 |
| CTA | 0.0525 ± 0.0039 | 0.0622 ± 0.0128 | 0.0097 |
| CTT | 0.0552 ± 0.0035 | 0.0569 ± 0.0128 | 0.0017 |
| TGA | $0.0500 \pm 0.0035$ | $0.0561 \pm 0.0128$ | 0.0062 |
| TGT | $0.0527 \pm 0.0039$ | $0.0478 \pm 0.0128$ | -0.0049 |
| TTT | $0.0500 \pm 0.0035$ | $0.0598 \pm 0.0128$ | 0.0098 |
| TTA | $0.0473 \pm 0.0013$ | $0.0155 \pm 0.0128$ | -0.0318 |

**Table S5.**
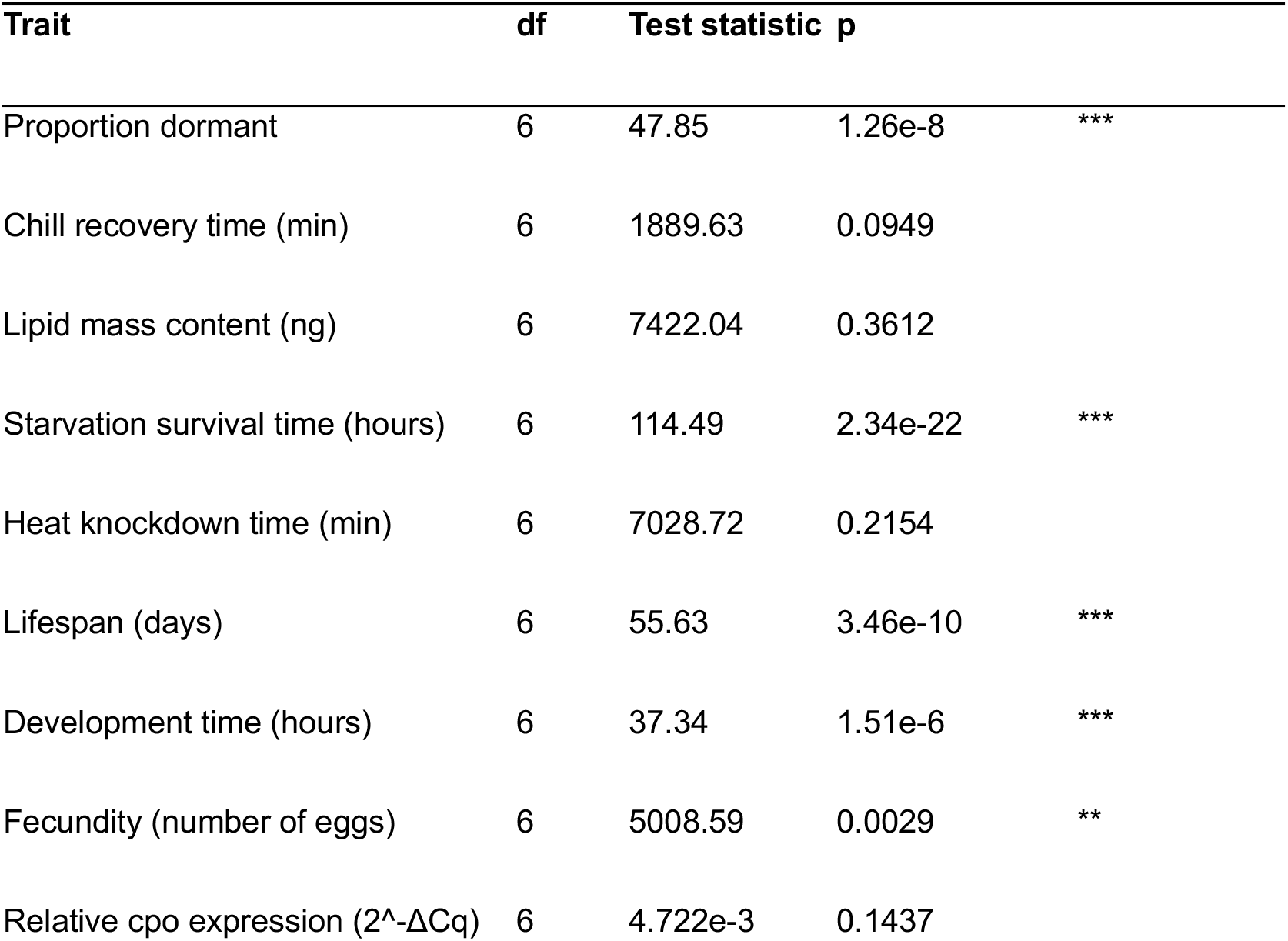
Likelihood-ratio test for the combined effect of three-locus genotype (Combination, 7 levels corresponding to the established ROPs) on each trait, comparing an intercept-only model to a model with Combination as a single categorical predictor. Reported as deviance for the binomial dormancy model, Chisq for traits with a bottle/vial random effect (starvation, lifespan, development), and a Sum-of-Squares-based F-test for all other continuous traits; see Methods. * p<0.05, ** p<0.01, *** p<0.001.

**Table S6.** Likelihood-ratio test for genotype-combination effects beyond additive main effects, comparing an additive model (Trait ∼ *cpo*^4408^ + *cpo*^5558^ + *cpo*^7866^) to the full three-locus Combination model, for each trait. This isolates the epistatic component specifically, controlling for additive main effects, and complements the vertex-specific deviation tests reported in Table S7. Because the *cpo*^CGA^ combination is absent from the data, individual 2-way and 3-way interaction terms cannot be separated from one another and this test reports their combined contribution only. *p<0.05, ** p<0.01, *** p<0.001.

| Trait | df | Test statistic | p |  |
| --- | --- | --- | --- | --- |
| Proportion dormant | 3 | 10.12 | 0.0176 | * |
| Chill recovery time (min) | 3 | 932.97 | 0.1491 |  |
| Lipid mass content (ng) | 3 | 620.45 | 0.9077 |  |
| Starvation survival time (hours) | 3 | 94.94 | 1.90e-20 | *** |
| Heat knockdown time (min) | 3 | 3191.48 | 0.2863 |  |
| Lifespan (days) | 3 | 4.79 | 0.1878 |  |
| Development time (hours) | 3 | 24.41 | 2.06e-5 | *** |
| Fecundity (number of eggs) | 3 | 3252.74 | 0.0048 | ** |
| Relative *cpo* expression (2<sup>-ΔCq</sup>) | 3 | 3.000e-3 | 0.1077 |  |

**Table S7.**
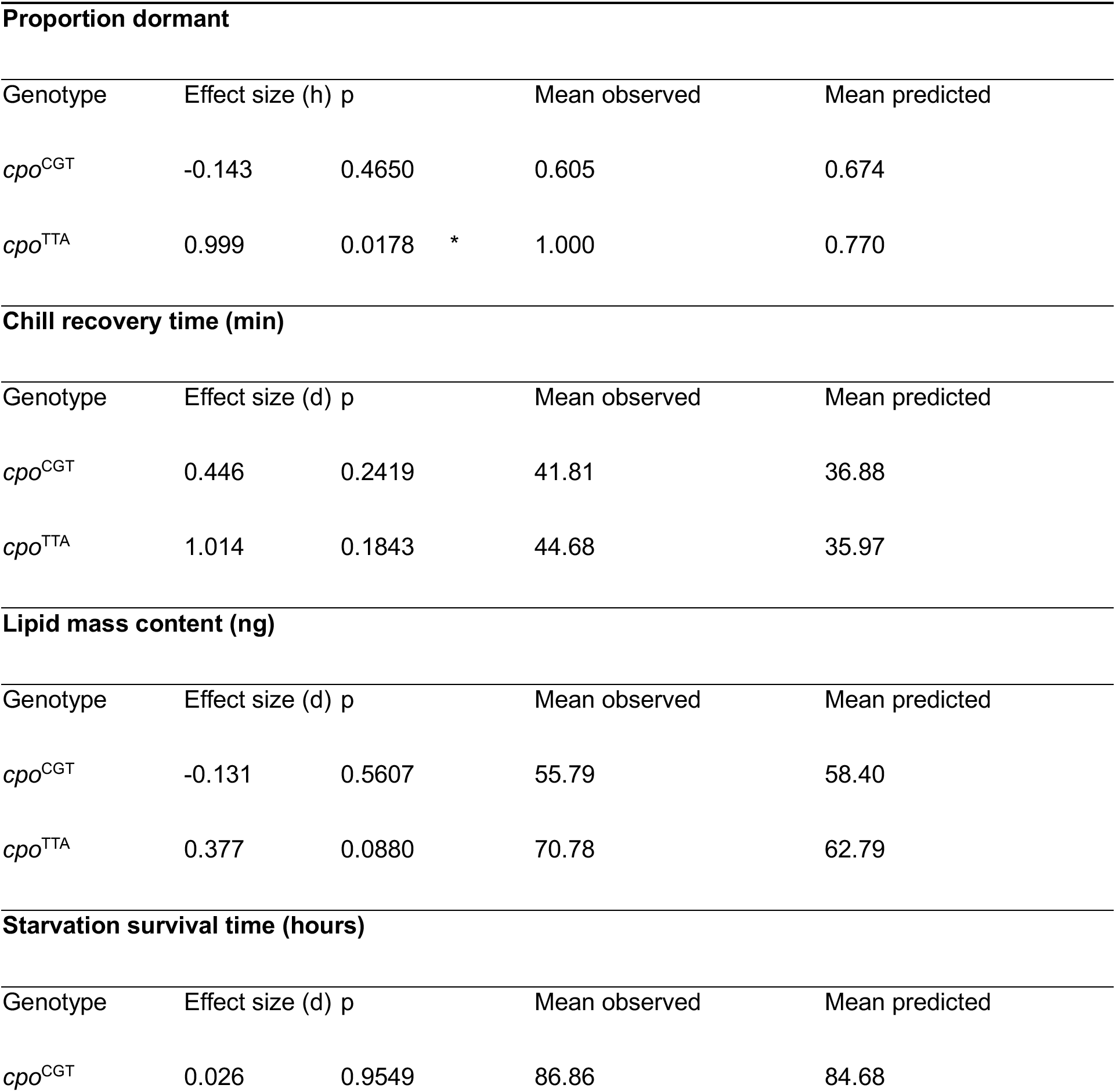
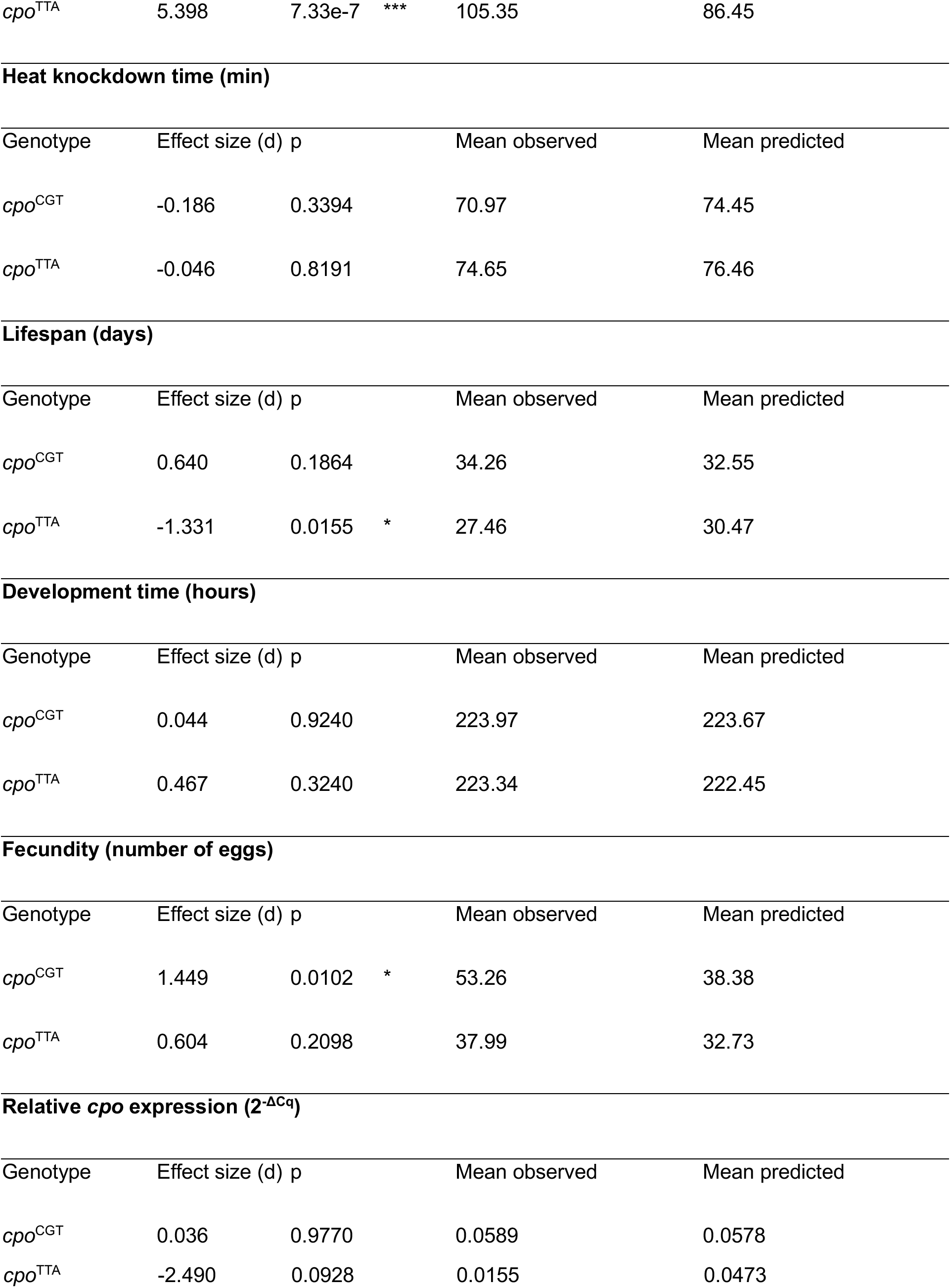
Deviation of observed trait values from additive null predictions for the tropical (*cpo*^CGT^) and temperate (*cpo*^TTA^) three-locus genotype combinations. For starvation resistance, lifespan, and development time, individual values were first averaged to bottle/vial-level means before testing (see Methods); for other continuous traits, individual values were compared directly. Effect sizes are Cohen’s h for the binomial dormancy proportion and Cohen’s d for all other traits. * p<0.05, ** p<0.01, *** p<0.001.

**Table S8.** Linkage disequilibrium (D’) among the three focal *couch potato* (*cpo*) loci in each wild-collected population, calculated directly from observed 3-locus haplotype counts (not the DGRP genotype-dosage approximation used in Methods/Figure 1). Note that D’ at small sample sizes (e.g. Florida, n=13) can be highly sensitive to individual haplotype counts and should be interpreted with caution.

| Population | Latitude | *cpo*<sup>4408</sup> – *cpo*<sup>5558</sup> | *cpo*<sup>5558</sup> – *cpo*<sup>7866</sup> | *cpo*<sup>4408</sup> – *cpo*<sup>7866</sup> |
| --- | --- | --- | --- | --- |
| Maine | 44.01°N | 0.181 | 0.439 | 0.350 |
| Pennsylvania | 39.12°N | 0.004 | 0.083 | 0.519 |
| North Carolina | 35.5°N | 0.136 | 0.035 | 0.548 |
| Florida | 25.5°N | 0.536 | 1.000 | 1.000 |

**Table S9.** Observed allele and three-locus genotype frequencies in wild-collected populations along the latitudinal cline, compared to predicted frequencies under random assortment of the three focal loci. Predicted genotype frequency is the product of the three single-locus allele frequencies observed in that population. Deviation is observed minus predicted.

| **Maine (N=79)** |  |  |  |  |
| --- | --- | --- | --- | --- |
| | n | Frequency ± SE | Predicted freq. | Deviation |
| *cpo*<sup>4408</sup> C | 27 | 0.342 ± 0.053 | — | — |
| *cpo*<sup>5558</sup> G | 29 | 0.367 ± 0.054 | — | — |
| *cpo*<sup>7866</sup> T | 34 | 0.430 ± 0.056 | — | — |
| *cpo*<sup>CGT</sup> | 5 | 0.063 ± 0.027 | 0.054 | 0.009 |
| *cpo*<sup>CGA</sup> | 8 | 0.101 ± 0.034 | 0.071 | 0.030 |
| *cpo*<sup>CTA</sup> | 2 | 0.025 ± 0.018 | 0.123 | -0.098 |
| *cpo*<sup>CTT</sup> | 12 | 0.152 ± 0.040 | 0.093 | 0.059 |
| *cpo*<sup>TGA</sup> | 14 | 0.177 ± 0.043 | 0.138 | 0.040 |
| *cpo*<sup>TGT</sup> | 2 | 0.025 ± 0.018 | 0.104 | -0.079 |
| *cpo*<sup>TTT</sup> | 15 | 0.190 ± 0.044 | 0.179 | 0.011 |
| *cpo*<sup>TTA</sup> | 21 | 0.266 ± 0.050 | 0.237 | 0.029 |
| **Pennsylvania (N=110)** |  |  |  |  |
| | n | Frequency ± SE | Predicted freq. | Deviation |
| *cpo*<sup>4408</sup> C | 52 | 0.473 ± 0.048 | — | — |
| *cpo*<sup>5558</sup> G | 40 | 0.364 ± 0.046 | — | — |
| *cpo*<sup>7866</sup> T | 66 | 0.600 ± 0.047 | — | — |
| *cpo*<sup>CGT</sup> | 16 | 0.145 ± 0.034 | 0.103 | 0.042 |
| *cpo*<sup>CGA</sup> | 3 | 0.027 ± 0.016 | 0.069 | -0.041 |
| *cpo*<sup>CTA</sup> | 7 | 0.064 ± 0.023 | 0.120 | -0.057 |
| *cpo*<sup>CTT</sup> | 26 | 0.236 ± 0.041 | 0.180 | 0.056 |
| *cpo*<sup>TGA</sup> | 15 | 0.136 ± 0.033 | 0.077 | 0.060 |
| *cpo*<sup>TGT</sup> | 6 | 0.055 ± 0.022 | 0.115 | -0.060 |
| *cpo*<sup>TTT</sup> | 18 | 0.164 ± 0.035 | 0.201 | -0.038 |
| *cpo*<sup>TTA</sup> | 19 | 0.173 ± 0.036 | 0.134 | 0.039 |

**North Carolina (DGRP) (N=145)**
|  | n | Frequency ± SE | Predicted freq. | Deviation |
| --- | --- | --- | --- | --- |
| *cpo*<sup>4408</sup> C | 72 | 0.497 ± 0.042 | — | — |
| *cpo*<sup>5558</sup> G | 46 | 0.317 ± 0.039 | — | — |
| *cpo*<sup>7866</sup> T | 96 | 0.662 ± 0.039 | — | — |
| *cpo*<sup>CGT</sup> | 24 | 0.166 ± 0.031 | 0.104 | 0.061 |
| *cpo*<sup>CGA</sup> | 2 | 0.014 ± 0.010 | 0.053 | -0.039 |
| *cpo*<sup>CTA</sup> | 9 | 0.062 ± 0.020 | 0.115 | -0.052 |
| *cpo*<sup>CTT</sup> | 37 | 0.255 ± 0.036 | 0.224 | 0.031 |
| *cpo*<sup>TGA</sup> | 13 | 0.090 ± 0.024 | 0.054 | 0.036 |
| $cpo^{TGT}$ | 7 | $0.048 \pm 0.018$ | 0.106 | -0.057 |
| $cpo^{TTT}$ | 28 | $0.193 \pm 0.033$ | 0.228 | -0.034 |
| $cpo^{TTA}$ | 25 | $0.172 \pm 0.031$ | 0.116 | 0.056 |

**Florida (Homestead) (N=13)**
| | n | Frequency ± SE | Predicted freq. | Deviation |
| --- | --- | --- | --- | --- |
| $cpo^{4408}$ C | 4 | $0.308 \pm 0.128$ | — | — |
| $cpo^{5558}$ G | 6 | $0.462 \pm 0.138$ | — | — |
| $cpo^{7866}$ T | 10 | $0.769 \pm 0.117$ | — | — |
| $cpo^{CGT}$ | 3 | $0.231 \pm 0.117$ | 0.109 | 0.122 |
| $cpo^{CGA}$ | 0 | $0.000 \pm 0.000$ | 0.033 | -0.033 |
| $cpo^{CTA}$ | 0 | $0.000 \pm 0.000$ | 0.038 | -0.038 |
| $cpo^{CTT}$ | 1 | $0.077 \pm 0.074$ | 0.127 | -0.051 |
| $cpo^{TGA}$ | 0 | $0.000 \pm 0.000$ | 0.074 | -0.074 |
| $cpo^{TGT}$ | 3 | $0.231 \pm 0.117$ | 0.246 | -0.015 |
| $cpo^{TTT}$ | 3 | $0.231 \pm 0.117$ | 0.287 | -0.056 |
| $cpo^{TTA}$ | 3 | $0.231 \pm 0.117$ | 0.086 | 0.145 |

**Table S10.** Observed allele and three-locus genotype frequencies across seasonal collections (May, June, October) in a Pennsylvania orchard, compared to predicted frequencies under random assortment of the three focal loci within each collection. Predicted genotype frequency is the product of the three single-locus allele frequencies observed in that collection. Deviation is observed minus predicted.

| **May (N=36)** |  |  |  |  |
| --- | --- | --- | --- | --- |
| | n | Frequency ± SE | Predicted freq. | Deviation |
| *cpo*<sup>4408</sup> C | 19 | 0.528 ± 0.083 | — | — |
| *cpo*<sup>5558</sup> G | 10 | 0.278 ± 0.075 | — | — |
| *cpo*<sup>7866</sup> T | 13 | 0.361 ± 0.080 | — | — |
| *cpo*<sup>CGT</sup> | 6 | 0.167 ± 0.062 | 0.094 | 0.073 |
| *cpo*<sup>CGA</sup> | 1 | 0.028 ± 0.027 | 0.053 | -0.025 |
| *cpo*<sup>CTA</sup> | 3 | 0.083 ± 0.046 | 0.138 | -0.054 |
| *cpo*<sup>CTT</sup> | 9 | 0.250 ± 0.072 | 0.244 | 0.006 |
| *cpo*<sup>TGA</sup> | 1 | 0.028 ± 0.027 | 0.047 | -0.020 |
| *cpo*<sup>TGT</sup> | 2 | 0.056 ± 0.038 | 0.084 | -0.028 |
| *cpo*<sup>TTT</sup> | 6 | 0.167 ± 0.062 | 0.218 | -0.051 |
| *cpo*<sup>TTA</sup> | 8 | 0.222 ± 0.069 | 0.123 | 0.099 |
| **June (N=42)** |  |  |  |  |
| | n | Frequency ± SE | Predicted freq. | Deviation |
| *cpo*<sup>4408</sup> C | 21 | 0.500 ± 0.077 | — | — |
| *cpo*<sup>5558</sup> G | 14 | 0.333 ± 0.073 | — | — |
| *cpo*<sup>7866</sup> T | 21 | 0.500 ± 0.077 | — | — |
| *cpo*<sup>CGT</sup> | 4 | 0.095 ± 0.045 | 0.083 | 0.012 |
| *cpo*<sup>CGA</sup> | 2 | 0.048 ± 0.033 | 0.083 | -0.036 |
| *cpo*<sup>CTA</sup> | 5 | 0.119 ± 0.050 | 0.167 | -0.048 |
| *cpo*<sup>CTT</sup> | 10 | 0.238 ± 0.066 | 0.167 | 0.071 |
| *cpo*<sup>TGA</sup> | 7 | 0.167 ± 0.058 | 0.083 | 0.083 |
| *cpo*<sup>TGT</sup> | 1 | 0.024 ± 0.024 | 0.083 | -0.060 |
| *cpo*<sup>TTT</sup> | 6 | 0.143 ± 0.054 | 0.167 | -0.024 |
| *cpo*<sup>TTA</sup> | 7 | 0.167 ± 0.058 | 0.167 | 0.000 |

**October (N=34)**
|  | n | Frequency ± SE | Predicted freq. | Deviation |
| --- | --- | --- | --- | --- |
| *cpo*<sup>4408</sup> C | 17 | 0.500 ± 0.086 | — | — |
| *cpo*<sup>5558</sup> G | 13 | 0.382 ± 0.083 | — | — |
| *cpo*<sup>7866</sup> T | 14 | 0.412 ± 0.084 | — | — |
| *cpo*<sup>CGT</sup> | 4 | 0.118 ± 0.055 | 0.112 | 0.005 |
| *cpo*<sup>CGA</sup> | 1 | 0.029 ± 0.029 | 0.079 | -0.049 |
| *cpo*<sup>CTA</sup> | 3 | 0.088 ± 0.049 | 0.127 | -0.039 |
| *cpo*<sup>CTT</sup> | 9 | 0.265 ± 0.076 | 0.182 | 0.083 |
| $cpo^{TGA}$ | 7 | $0.206 \pm 0.069$ | 0.079 | 0.127 |
| $cpo^{TGT}$ | 1 | $0.029 \pm 0.029$ | 0.112 | -0.083 |
| $cpo^{TTT}$ | 6 | $0.176 \pm 0.065$ | 0.182 | -0.005 |
| $cpo^{TTA}$ | 3 | $0.088 \pm 0.049$ | 0.127 | -0.039 |

**Figure S1.** Schematic of the recombinant outbred population (ROP) design and additive null prediction model. DGRP founder lines sharing the same three-locus *cpo* genotype (e.g., *cpo*^TTA^ or *cpo*^CGT^, shown) were crossed to generate genetically heterogeneous ROPs fixed only at the three focal SNPs (*cpo*^4408^, *cpo*^5558^, *cpo*^7866^), controlling for background genetic variation while holding the focal combination constant. Seven of the eight possible three-locus combinations were established as ROPs (*cpo*^CGA^ was too rare in the DGRP to identify sufficient founders). The additive null prediction for a given combination (e.g., *cpo*^TTA^) was calculated by averaging the marginal effect of each focal allele across all genotype classes carrying it, then combining the three single-locus marginal effects; deviation of the observed ROP phenotype from this additive prediction constitutes the epistasis test used throughout Figures 2–3.

**Figure S2.** Validation of ROP phenotypes in independently-derived Pennsylvania inbred lines fixed for the tropical (*cpo*^CGT^) and temperate (*cpo*^TTA^) three-locus combinations. Light bars show the additive null prediction (mean of marginal single-locus effects); dark bars show the observed value. Significance brackets indicate within-genotype deviation from the additive prediction (t-test) and the cross-genotype comparison (*cpo*^CGT^ vs. *cpo*^TTA^, observed values); effect sizes are Cohen’s d. (A) Chill coma recovery time. (B) Starvation survival. (C) Heat knockdown time. Error bars show ±1 SE. Effect sizes here are substantially smaller than in the corresponding ROP comparisons (Figure 2), consistent with the inbred lines testing within-line phenotypic consistency on a distinct genetic background rather than replicating the full compositional epistasis design.

**Figure S3.** Robustness of *EcR::usp* and *cnc::maf-S* enrichment relative to background *cpo* intronic variation. (A) Total disrupted motifs per SNP (|score difference| > 0.1) for the three focal clinal SNPs, 41 additional clinal SNPs from the same region, and 30 non-clinal SNPs from the same intronic window; groups do not differ significantly (Kruskal-Wallis p = 0.783). Focal SNPs are individually marked (triangle = *cpo*4408, circle = *cpo*5558, square = *cpo*7866). (B) Proportion of SNPs disrupting *EcR::usp* or *cnc::maf-S*, focal (n=3) versus all non-focal SNPs combined (n=71); focal SNPs are significantly enriched for both motifs (Fisher’s exact p = 0.029 and p = 0.011, respectively). (C) Clinal significance (−log10 CQ) for clinal SNPs (focal + other-clinal) split by hit status for each motif; neither *EcR::usp* (Wilcoxon p = 0.197) nor *cnc::maf-S* (Wilcoxon p = 0.029) shows a strong universal association with clinal significance in isolation. (D) Extending the comparison across all 105 testable JASPAR motifs (motifs with ≥2 disrupting and ≥2 non-disrupting SNPs among the 44 clinal SNPs), ranked by Wilcoxon p-value; point size indicates the number of disrupting SNPs supporting each test. cnc::maf-S ranks 3^rd^ of 105; the two motifs ranking above it are each supported by only 2 disrupting SNPs (the minimum testable sample size in this analysis) and are not interpreted as stronger competing signals given this limited support. *EcR::usp* does not reach significance in this motif-wide framework despite having the largest sample size (n=8) among top-ranked motifs, though the direction of effect is consistent with the other results (higher clinal significance among disrupting SNPs). Dashed line marks p = 0.05.

**Figure S4.** Combined neighbor-joining tree (unrooted) of the *couch potato* (*cpo*) focal region, with 2,897 shared SNPs after liftover across n=338 samples from all three sampled populations: Maine (triangle), Pennsylvania (spring=circle, autumn=square), North Carolina (diamond). Tip and branch color indicate three-locus haplotype. Branch lengths reflect genetic distance across the shared SNP set only and should not be interpreted as absolute divergence.

