## Supplemental Figures for "Intragenic epistasis in couch potato shapes life history traits and climatic adaptation in *Drosophila melanogaster*"

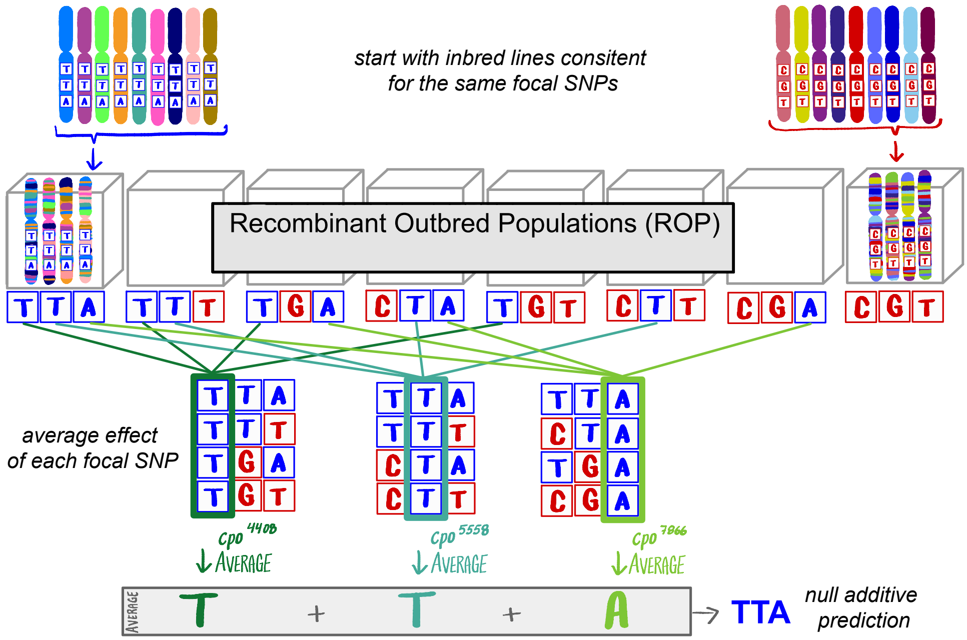


**Figure S1.** Schematic of the recombinant outbred population (ROP) design and additive null prediction model. DGRP founder lines sharing the same three-locus *cpo* genotype (e.g., TTA or CGT, shown) were crossed to generate genetically heterogeneous ROPs fixed only at the three focal SNPs (*cpo*$^{4408}$, *cpo*$^{5558}$, *cpo*$^{7866}$), controlling for background genetic variation while holding the focal combination constant. Seven of the eight possible three-locus combinations were established as ROPs (*cpo*$^{CGA}$ was too rare in the DGRP to identify sufficient founders). The additive null prediction for a given combination (e.g., *cpo*$^{TTA}$) was calculated by averaging the marginal effect of each focal allele across all genotype classes carrying it, then combining the three single-locus marginal effects; deviation of the observed ROP phenotype from this additive prediction constitutes the epistasis test used throughout Figures 2–3.

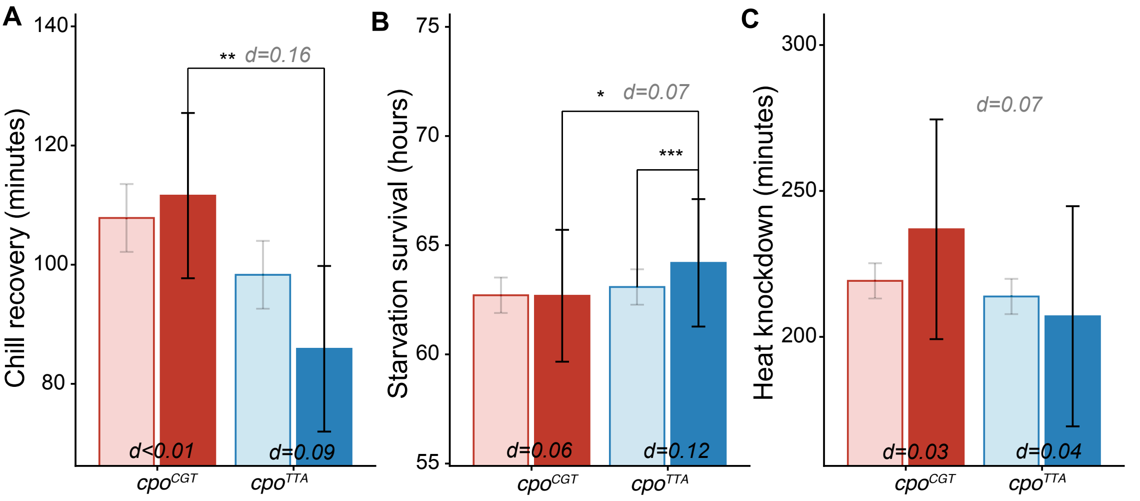


**Figure S2.** Validation of ROP phenotypes in independently-derived Pennsylvania inbred lines fixed for the tropical (*cpo*$^{CGT}$) and temperate (*cpo*$^{TTA}$) three-locus combinations. Light bars show the additive null prediction (mean of marginal single-locus effects); dark bars show the observed value. Significance brackets indicate within-genotype deviation from the additive prediction (t-test) and the cross-genotype comparison (CGT vs. TTA, observed values); effect sizes are Cohen's d. (A) Chill coma recovery time. (B) Starvation survival. (C) Heat knockdown time. Error bars show ±1 SE. Effect sizes here are substantially smaller than in the corresponding ROP comparisons (Figure 2), consistent with the inbred lines testing within-line phenotypic consistency on a distinct genetic background rather than replicating the full compositional epistasis design.

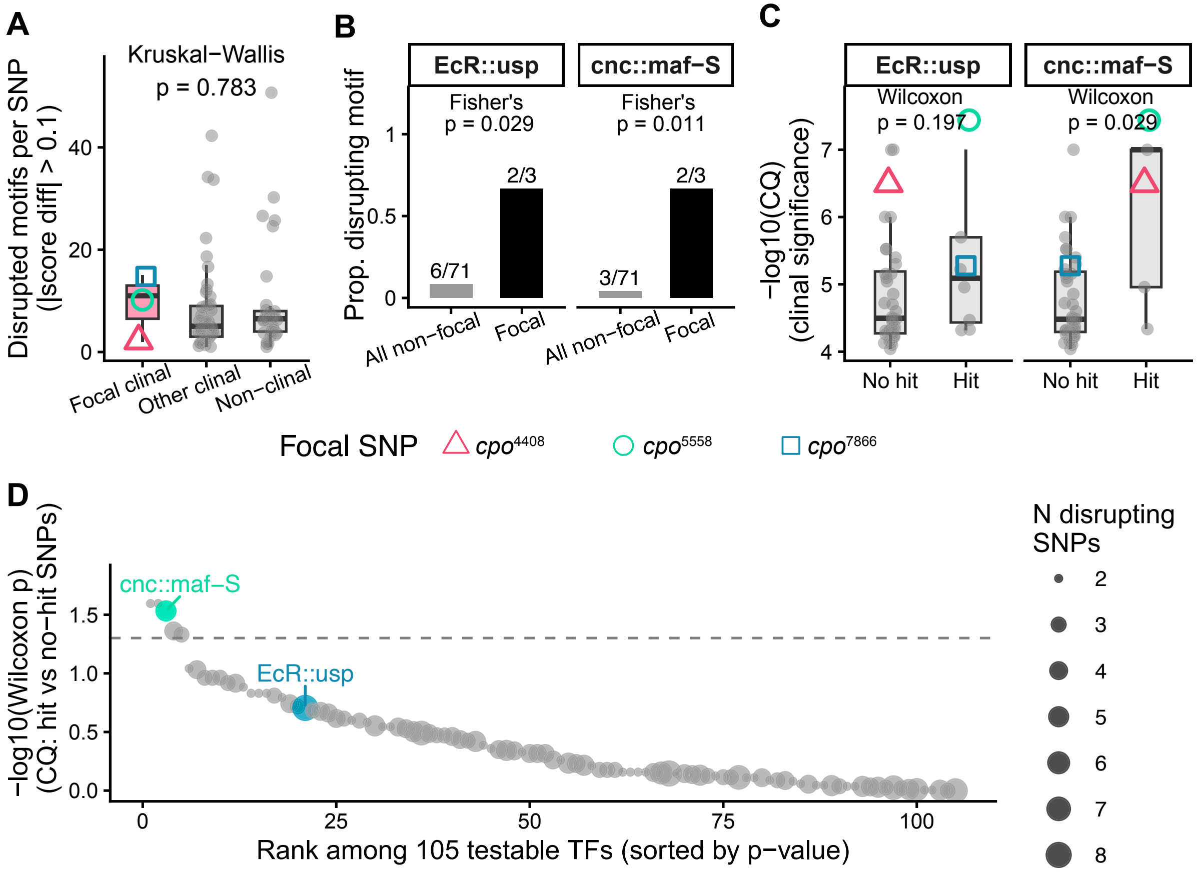


**Figure S3.** Robustness of *EcR::usp* and *cnc::maf-S* enrichment relative to background *cpo* intronic variation. (A) Total disrupted motifs per SNP (|score difference| > 0.1) for the three focal clinal SNPs, 41 additional clinal SNPs from the same region, and 30 non-clinal SNPs from the same intronic window; groups do not differ significantly (Kruskal-Wallis p = 0.783). Focal SNPs are individually marked (triangle = *cpo*$^{4408}$, circle = *cpo*$^{5558}$, square = *cpo*$^{7866}$). (B) Proportion of SNPs disrupting *EcR::usp* or *cnc::maf-S*, focal (n=3) versus all non-focal SNPs combined (n=71); focal SNPs are significantly enriched for both motifs (Fisher's exact p = 0.029 and p = 0.011, respectively). (C) Clinal significance (−log10 CQ) for clinal SNPs (focal + other-clinal) split by hit status for each motif; neither *EcR::usp* (Wilcoxon p = 0.197) nor *cnc::maf-S* (Wilcoxon p = 0.029) shows a strong universal association with clinal significance in isolation. (D) Extending the comparison across all 105 testable JASPAR motifs (motifs with ≥2 disrupting and ≥2 non-disrupting SNPs among the 44 clinal SNPs), ranked by Wilcoxon p-value; point size indicates the number of disrupting SNPs supporting each test. cnc::maf-S ranks 3rd of 105; the two motifs ranking above it are each supported by only 2 disrupting SNPs (the minimum testable sample size in this analysis) and are not interpreted as stronger competing signals given this limited support. *EcR::usp* does not reach significance in this motif-wide framework despite having the largest sample size (n=8) among top-ranked motifs, though the direction of effect is consistent with the other results (higher clinal significance among disrupting SNPs). Dashed line marks p = 0.05.

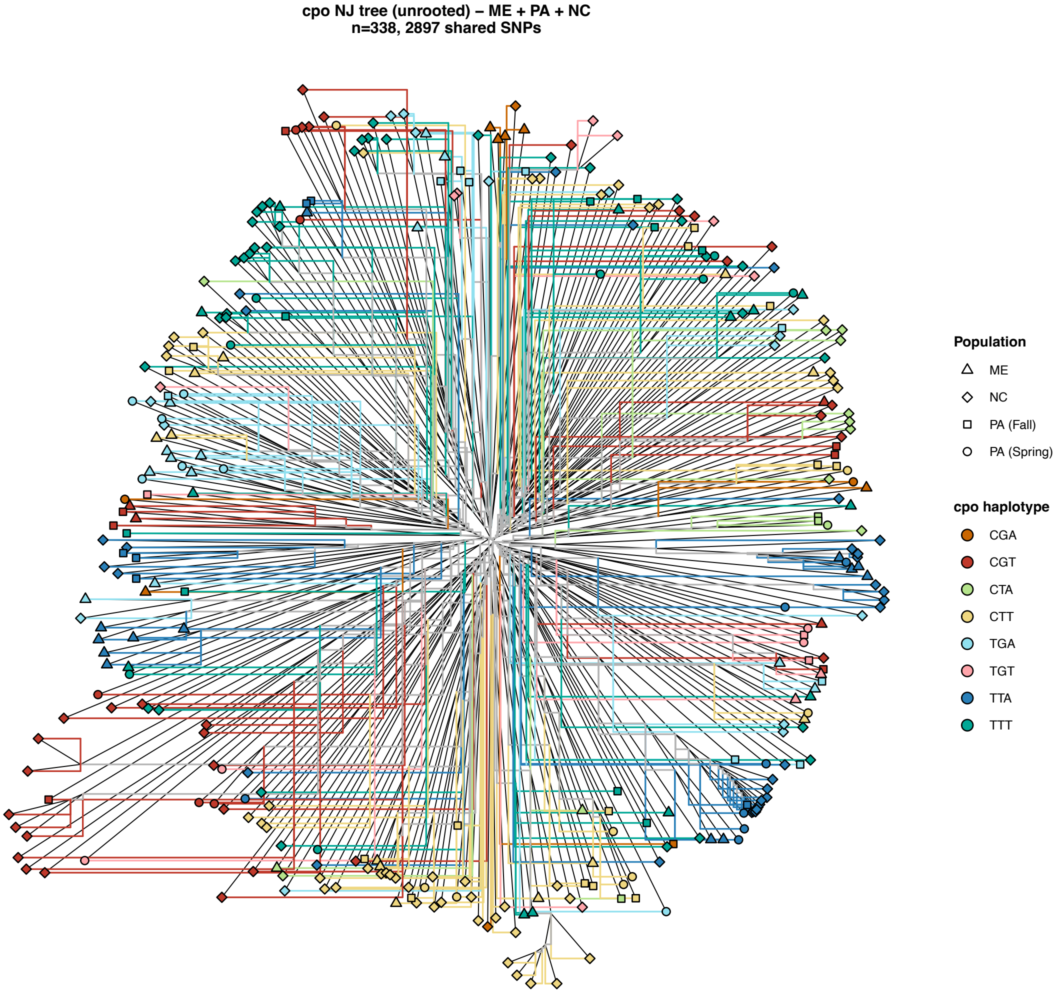


**Figure S4.** Combined neighbor-joining tree (unrooted) of the *couch potato* (*cpo*) focal region, with 2,897 shared SNPs after liftover across n=338 samples from all three sampled populations: Maine (triangle), Pennsylvania (spring=circle, autumn=square), North Carolina (diamond). Tip and branch color indicate three-locus haplotype. Branch lengths reflect genetic distance across the shared SNP set only and should not be interpreted as absolute divergence.
